# Syngap1 Synchronizes Relative Neuronal Maturation Across Cortical Areas to Organize Distributed Functional Networks

**DOI:** 10.64898/2026.03.27.714839

**Authors:** Randall M. Golovin, Bianca Garcia-Gonzalez, Sheldon D. Michaelson, Massimiliano Aceti, Seth Butz, Camilo Rojas, Courtney A. Miller, Thomas Vaissiere, Gavin Rumbaugh

**Affiliations:** Departments of Neuroscience and Molecular Medicine, The Herbert Wertheim UF Scripps Institute for Biomedical Innovation & Technology, Jupiter, FL, USA; The Skaggs Graduate School of Chemical and Biological Sciences at Scripps Research, Jupiter, FL, USA; Department of Drug Discovery, H. Lee Moffitt Cancer Center and Research Institute, Tampa, FL

## Abstract

Neurodevelopmental disorders are increasingly viewed as disorders of distributed brain networks, yet how developmental perturbations generate co-occurring hypo- and hyper-functional network states remains unclear. We show that *Syngap1* haploinsufficiency in mice produces opposing cortical activity patterns: sensory-evoked responses exhibit reduced gain, whereas state-linked movement/arousal activity is amplified. Restricting *Syngap1* deficiency to developing cortical excitatory neurons reproduces distributed sensory hypofunction but not movement-linked hyperfunction, indicating non-uniform, cell-type-dependent contributions to these patterns. During a postnatal window of circuit assembly, Layer 2/3 intratelencephalic populations across cortical regions occupy distinct maturation states, reflected in separable dendritic architectures and intrinsic excitability. Perturbing *Syngap1* reduces this developmental separability through region-dependent, opposite-direction effects on dendritic maturation and inversion of ERK-sensitive control of intrinsic excitability. We propose that coordinated relative maturation of interacting neuronal populations establishes distributed network balance, and that gene-dependent shifts in this coordination produce stable, opposing network states.

## INTRODUCTION

Neurodevelopmental disorders (NDDs), including autism spectrum disorder (ASD) and intellectual disability (ID), are characterized by impairments in cognition, communication, and behavioral flexibility. In addition to these core features, atypical sensory processing, motor control, and arousal regulation are highly prevalent and clinically significant [1, 2]. These domains are normally tightly integrated to instruct adaptive behavior: sensory inputs are interpreted in the context of behavioral state to guide action, supported by widespread, state-dependent modulation of cortical activity [3–6].

Despite this integration, large-scale neuroimaging studies have revealed a persistent organizational paradox in ASD. Rather than uniform increases or decreases in connectivity or function, ASD is associated with spatially structured mosaics of network activity, including coexisting hypo- and hyper-connectivity across cortical systems [7–10]. Sensory and sensorimotor networks often show reduced functional coupling, whereas higher-order association networks can be relatively amplified depending on brain state and measurement modality [11, 12]. These findings indicate that network dysfunction in ASD is not uniform, but instead organized into opposing functional network states within the same brain. These opposing network states are associated with symptom severity, consistent with the idea that systems-level imbalances contribute to core behavioral features of ASD.

A central unresolved question is how such structured, bidirectional network states arise during development. Influential models centered on excitation/inhibition imbalance or synaptic plasticity have provided important insight into local circuit dysfunction [13, 14]. However, these models do not readily explain how opposing alterations emerge across distributed subnetworks. This limitation highlights the need for biological frameworks that operate at the level of distributed networks rather than local circuit properties.

One potential resolution is that neurodevelopmental perturbations alter the relative timing of neuronal maturation across interacting circuits. Developmental neuroscience has established that the timing and rate of maturation constrain synaptic refinement and circuit assembly, including through critical periods of experience-dependent plasticity. If interacting cortical populations mature at different rates, their relative developmental positioning may shape how signals are integrated across networks. Differences in maturation state across neuronal populations in different cortical regions could therefore produce a developmental mosaic, in which populations occupy distinct positions along their maturation trajectories. Shifts in these relative positions could bias the balance between sensory-driven and state-dependent activity. Consistent with this view, cortical circuits mature along sensory–associational and anterior–posterior gradients [15], supported by regionally patterned gene-expression programs [16], and exhibit continuous gradients of functional organization [17]. However, whether developmental perturbations disrupt coordination across these interacting populations, and how this scales to mesoscale network dynamics, remains unknown.

Genetic studies show that many NDD risk genes contribute to shared liability across neurodevelopmental and psychiatric conditions [18–21], raising the possibility that gene-level disruptions during development bias distributed network architecture. Among these, *SYNGAP1* provides a powerful entry point. Haploinsufficiency of *SYNGAP1* in humans causes developmental encephalopathy characterized by intellectual disability, epilepsy, and autistic features [22, 23]. In rodents, *Syngap1* deficiency disrupts sensory processing, impairs sensorimotor dynamics, and produces cognitive and behavioral abnormalities) [24–26]. At the synaptic and cellular levels, *Syngap1* regulates dendritic growth, synapse maturation, intrinsic excitability, and critical-period plasticity [27–30]. Importantly, these effects are not uniform across cell types or regions, and SynGAP isoforms exhibit spatially and temporally distinct expression patterns [31, 32]. Together, these observations suggest that *Syngap1* does not act as a uniform regulator of neuronal function, but instead contributes to region- and cell-type-specific maturation programs that organize distributed neural systems.

Here, we combine cell- and region-specific genetic manipulations with measurements of neuronal morphology, intrinsic excitability, mesoscale cortical activity, and behavior in awake mice to determine how *Syngap1* shapes distributed cortical network organization. We show that *Syngap1* haploinsufficiency produces coexisting, opposing activity states within the same animal model: sensory-evoked responses exhibit reduced gain, whereas movement/state-linked activity is amplified. Restricting *Syngap1* deficiency to cortical excitatory neurons reproduces sensory hypofunction but not movement-linked hyperfunction, revealing non-uniform, cell-type-dependent contributions to these network states. During a defined postnatal window of circuit assembly, *Syngap1* exerts region-dependent, pleiotropic effects on layer 2/3 (L2/3) intratelencephalic (IT) neurons across connected cortical regions. Dendritic maturation is regulated cell-autonomously in opposite directions, whereas intrinsic excitability landscapes diverge across regions and depend on circuit context, with genotype-dependent inversion of Erk-sensitive signaling. Together, these findings demonstrate that perturbation of a single NDD risk gene reduces the separability of maturation states across interacting neuronal populations. We propose that coordinated relative maturation of distributed cortical ensembles establishes network balance, and that disruption of this coordination provides a developmental basis for the emergence of stable, opposing network states.

## RESULTS

### Syngap1 deficiency reduces distributed sensory-evoked cortical response gain

Prior studies have shown that *Syngap1* haploinsufficiency weakens whisker-evoked sensory responses within primary somatosensory cortex [30], consistent with deficits in both passive and active tactile processing [26]. However, it remains unknown to what extent other sensory modalities are affected, how sensory signals propagate as they flow to other cortical regions, and how gene deficiency at the cellular level may contribute to distributed sensory coding. To address these questions, we measured sensory-evoked cortical activity across the dorsal surface of the brain in awake head-fixed mice using widefield GCaMP6 imaging (*hemodynamically corrected ΔF/F*; **Fig. 1A)**. Because spontaneous movements strongly modulate cortical activity and can obscure sensory-evoked signals [4], analyses were restricted to trials without movement immediately before and after stimulus presentation.

**Figure 1.**
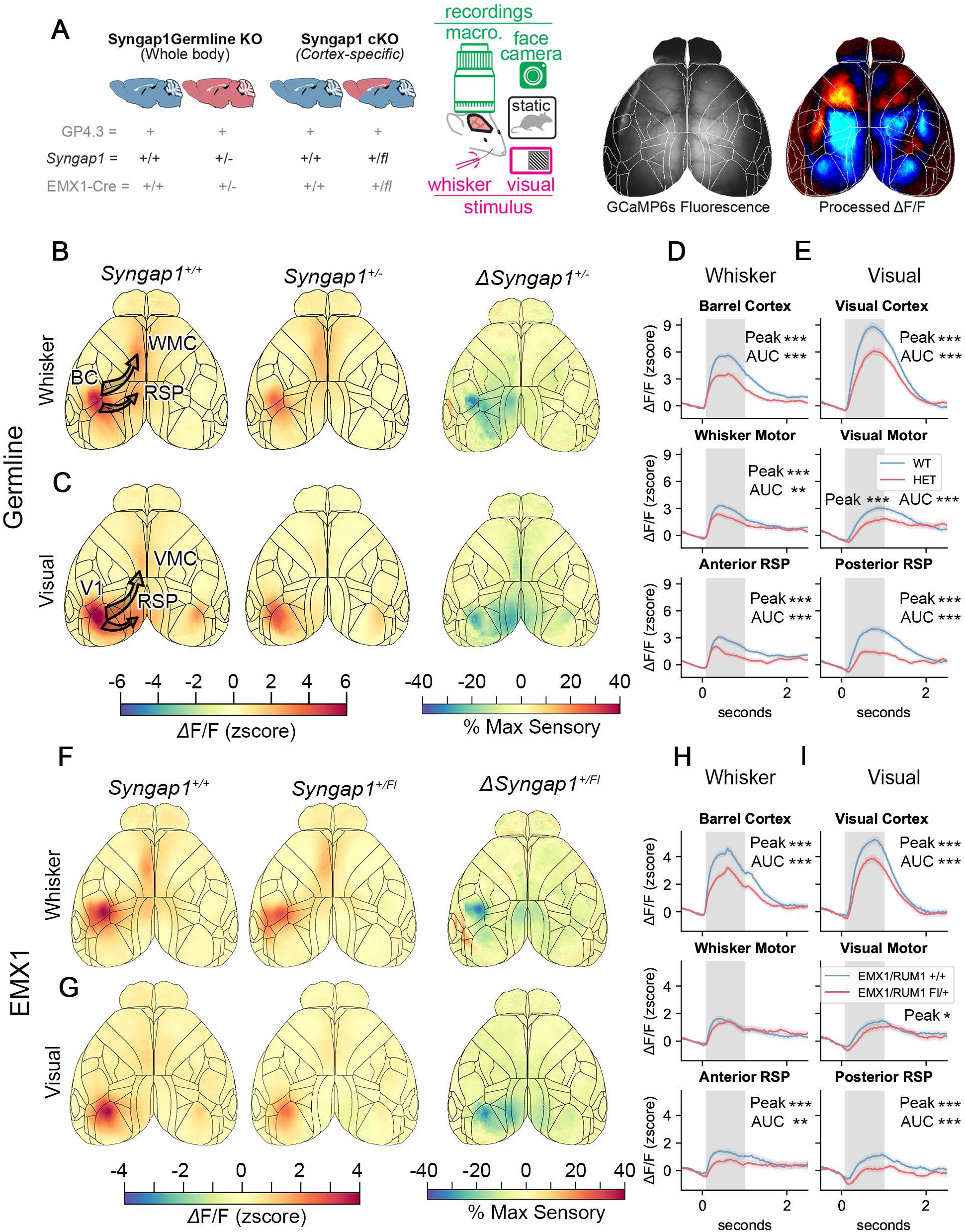
Syngap1 deficiency is associated with distributed sensory hypofunction that is recapitulated by cortex-restricted deletion. **(A)** Experimental schematic illustrating widefield GCaMP6 imaging of dorsal cortex in awake, head-fixed mice during controlled sensory stimulation. Hemodynamically corrected b..F/F signals were analyzed during movement-free trials to isolate stimulus-evoked activity. **(B-E)** Sensory-evoked cortical responses in germline Syngap1 heterozygous mice. Time-compressed b..F/F dorsal cortex maps (B) and time series across selected cortical regions (D) in response to whisker stimulation (Syngap1+/+: N=753 trials, N=7 mice, Syngap1+/-: N=462 trials, N=S mice). Similar maps (C) and regional time series (E) were generated in response to visual stimulation (Syngap1+/+: N=502 trials, N=6 mice, Syngap1+/-: N=454 trials, N=6 mice). **(F-I)** Maps and time series for either whisker (Syngap1+/+: N=584 trials, N=8 mice, Syngap1+/fl: N=637 trials, N=8 mice) or visual stimuli (Syngap1+/+: N=696 trials, N=7 mice, Syngap1+/fl: N=636 trials, N=7 mice) in cortical excitatory neuron-restricted Emx1-Cre Syngap1 conditional heterozygous mice. Brain maps (B,C,F,G) show the mean signal averaged across trials from 0.1-1 s post stimulus onset (left, shaded region of traces), or the difference between mutant and control averages expressed as a percentage of the max activity within barrel cortex (BC) or visual cortex (V1) in control animals. Traces show the mean ± SEM of small 10 x 10 pixel regions of interest manually placed on the initial center of activity in the indicated regions. Statistical comparisons are of the AUC from 0.1-1s post stimulus (shaded region) between genotypes within each of the three regions for each strain and stimulus modality. Significance was assessed using a Mann-Whitney U test with a Bonferroni correction for multiple comparisons. Data are mean± SEM. _*,_ p < 0.05; _**,_ p < 0.01, _***_ p < 0.001.

In germline *Syngap1* haploinsufficient mice (*Syngap1^+/-^*mice, or germline *Syngap1* heterozygotes), whisker stimulation evoked significantly reduced response amplitudes in primary somatosensory cortex (S1) compared to wild-type littermates **(Fig. 1B,D; *Fig. S1A*)**, consistent with prior reports using complementary approaches [26, 30]. A comparable reduction in response amplitude was observed in primary visual cortex (V1) following visual stimulation **(Fig. 1C,E; *Fig. S1B*)**, indicating that sensory hypofunction generalizes beyond the whisker system and is not modality-specific. Reduced response amplitudes were also evident during multimodal whisker-visual stimulation **(Fig. S2)**, supporting a general reduction in sensory-evoked cortical responses rather than a modality-restricted deficit.

Because widefield imaging enables simultaneous measurement across cortical regions engaged during sensory processing, we next examined whether reduced sensory-evoked responses were confined to primary sensory cortex or extended across broader cortical networks. In *Syngap1^+/-^* mice, whisker-evoked activity was attenuated not only within S1, but also across several downstream cortical regions typically coactivated during tactile processing, including motor, medial-frontal, and retrosplenial areas **(Fig. 1D; *Fig. S1A*).** We also measured distributed signals across dorsal cortex brain regions using alternative approaches, including a threshold-based mask created from the germline wild-type signals **(Fig. S3A,B,D)**, as well as applying ROIs based on the Allen common coordinate framework (CCF) **(Fig. S4).** These alternative approaches resulted in similar differences between genotypes and strains. A similar set of findings was also observed for visual stimulations **(Fig. 1C-E; *Fig. S1B; Fig. S3F,G,I; Fig. S4*)**. Thus, *Syngap1* haploinsufficiency reduces sensory-evoked responses across distributed cortical networks, rather than being limited to primary sensory areas. Throughout, we use ‘sensory hypofunction’ to refer to reduced population-level response amplitude (gain), not altered tuning or selectivity; because widefield ΔF/F reports population-level activity, these data do not directly assay pathway-specific synaptic routing, which would require targeted circuit-level manipulations.

To determine whether impaired sensory network function reflects cortex-intrinsic mechanisms or secondary effects of broader circuit dysfunction, we performed widefield imaging experiments in *Emx1*-Cre *Syngap1* conditional heterozygous mice, in which *Syngap1* haploinsufficiency is selectively restricted to cortical excitatory projection neurons **(Fig. 1A)**. Importantly, *Emx1*-Cre is active during embryonic cortical development, enabling selective perturbation of *Syngap1* in cortical excitatory neurons throughout circuit assembly and early maturation. In this model, whisker- or visual-evoked responses in both S1 and V1 were similarly reduced relative to control littermates **(Fig. 1F,G)**, closely phenocopying the deficits observed in germline mutants. Attenuated sensory-evoked activity was likewise observed across multiple downstream cortical regions, including retrosplenial cortex and associative areas **(Fig. 1H,I; Fig. S1C-D; Fig. S3C,E, H,J)**. Notably, in contrast to the germline model, movement-free sensory stimulation produced minimal differences in motor cortical response amplitude in *Emx1*-Cre *Syngap1* mutants relative to controls **(Fig. 1H,I; Fig. S3C,E,H,J; Fig. S4)**. Together, these findings indicate that reduced sensory responses arise, at least in part, from *Syngap1* loss in cortical excitatory neurons. Moreover, the relative preservation of motor cortical responses in the cortex-restricted model suggests that sensory and motor network phenotypes in *Syngap1* deficiency are supported by partially distinct cellular and circuit mechanisms. Finally, these experiments directly demonstrate a reduction in sensory-evoked response amplitude across cortex; altered propagation across networks is inferred from the distributed reduction in response amplitude (*e.g., reduced amplitude in cortical regions known to be directly downstream of, and synaptically coupled to, primary cortex*).

Because distributed sensory hypofunction persists when *Syngap1* loss is restricted to cortical excitatory neurons, it is unlikely to arise solely from altered subcortical input or global state changes. These results are consistent with a cortex-intrinsic contribution to reduced sensory response gain. We therefore examined development of L2/3 IT neurons in primary sensory cortex as a candidate cellular substrate for reduced sensory response gain in *Syngap1* mice. These neurons receive substantial sensory input and provide long-range corticocortical projections to downstream cortical areas. We chose to focus on late postnatal development ∼(PND14-21) of these neurons for two reasons. First, this developmental window represents a critical period in which dendritic growth, synaptic refinement, and intrinsic excitability changes shape how neurons are dynamically integrated into cortical circuits [33, 34]. Second, *Syngap1* influences critical periods of glutamatergic sensory cortex neuronal maturation by regulating synapse dynamics essential for circuit assembly and refinement [27–29]. In addition, prior work has shown that adult *Syngap1* mutant mice exhibit reduced feedforward excitation in somatosensory cortex, driven by fewer excitatory synaptic connections [30]. Whether these circuit-level deficits arise during development remains unknown.

To assess dendritic morphogenesis in developing neurons *in vivo*, we crossed a *td*Tomato Cre reporter line with *Syngap1* germline mutant mice, with offspring injected with a low titer of AAV-cre to induce sparse recombination of the reporter throughout the brain **(Fig. 2A)**. We found that in germline *Syngap1* haploinsufficient mice, L2/3 glutamatergic neurons in both S1 and V1 exhibited shorter dendritic arbors and reduced complexity during postnatal development **(Fig. 2B,C)**. Moreover, in developing L2/3 neurons from *Syngap1* Hets, we also observed reduced dendritic spine density **(Fig. S5A)**. Reduced dendritic length together with decreased spine density is expected to reduce the total number of excitatory synapses. This was experimentally validated by counting dendritic spine synapses in upper lamina of S1 **(Fig. S5B).** Thus, critical sensory relay neurons in *Syngap1* mutants exhibit reduced overall anatomical excitatory synaptic input in this early stage of development. Furthermore, intrinsic neuronal excitability in L2/3 sensory neurons was also reduced relative to WT neurons **(Fig. 2D,E)**, an orthogonal finding that is also consistent with reduced recruitment of neurons into developing sensory circuits. Notably, slowed maturation at the morphological level was specific to upper lamina pyramidal neurons because basal dendrites from L5 neurons were longer in *Syngap1* deficient mice compared to wildtype littermates **(Fig. 2F),** a finding that agrees with past reports that measured the maturation of apical dendrites from L5b bi-tufted neurons in somatosensory cortex [27]. Thus, *Syngap1* deficiency selectively impacts upper-layer neuronal maturation in sensory cortex consistent with previously reported circuit deficits in the same area in adult mice.

**Figure 2.**
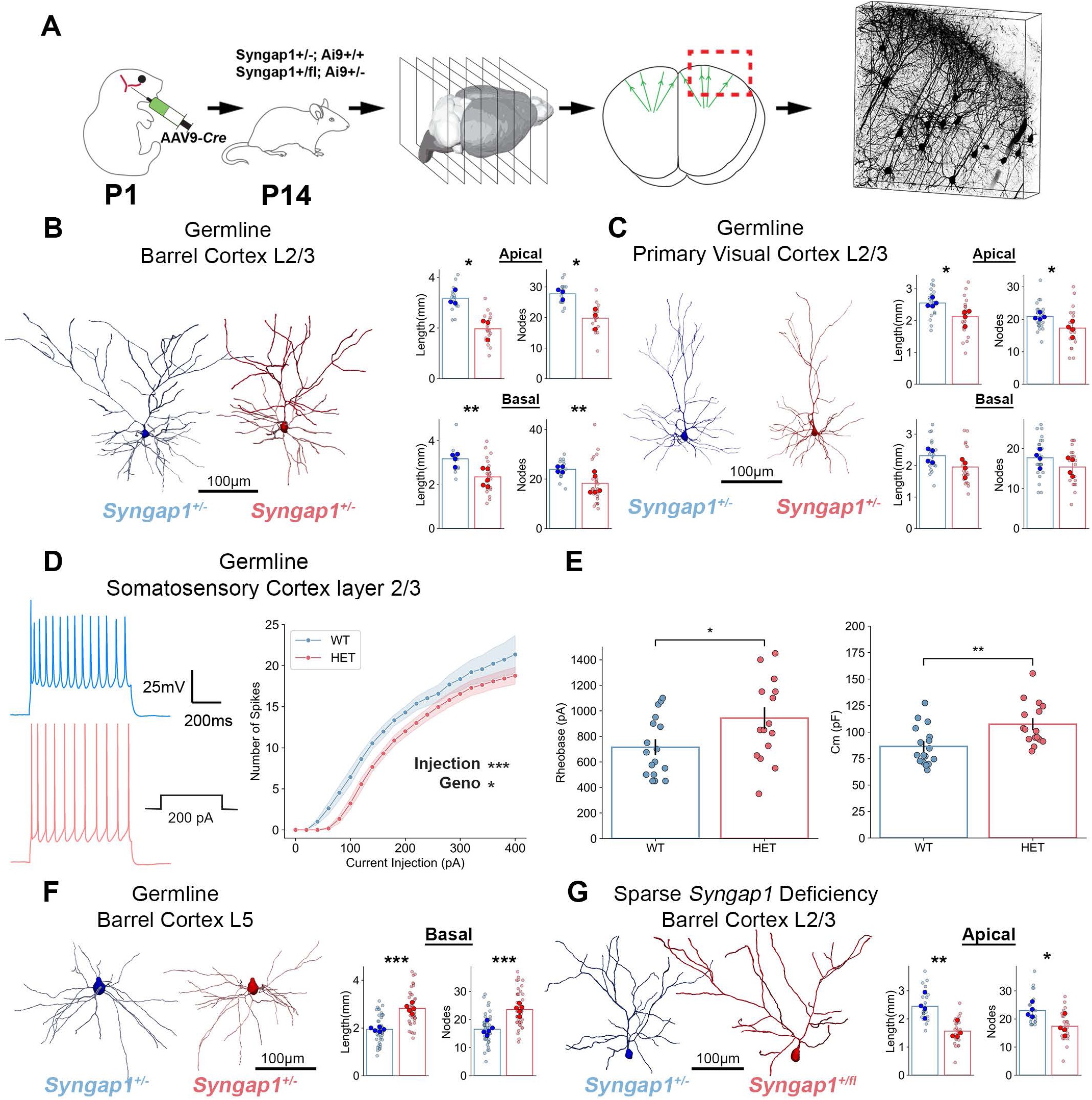
Syngap1 haploinsufficiency reduces dendritic maturation and intrinsic responsiveness of sensory L2/3 pyramidal neurons. **(A)** Experimental strategy for sparse labeling and reconstruction of L2/3 pyramidal neurons. **A** ere-dependent fluorescent reporter was sparsely activated using low-titer AAV-Cre to enable morphological analysis of individual neurons. (right) a representative image of sparse TdTomato expression within a S00µm thick section **(B-C)** Dendritic reconstructions and quantification of L2/3 pyramidal neurons in primary somatosensory (barrel) cortex (S1) and primary visual cortex (V1) from PND14 germline Syngap1 heterozygous mice and wild-type littermates. Syngap1 deficiency reduces total dendritic length and branch complexity in sensory cortex populations. S1 Apical: Syngap1+/+ N=15 neurons, N=3 mice, Syngap1+-+ N=18 neurons, N=3 mice. S1 Basal: Syngap1+/+ N=15 neurons, N=4 mice, Syngap1+-+ N=23 neurons, N=S mice. V1 Apical: Syngap1+/+ N=24 neurons, N=4 mice, Syngap1+-+ N=23 neurons, N=4 mice. V1 Basal: Syngap1+/+ N=24 neurons, N=4 mice, Syngap1+-+ N=23 neurons, N=4 mice. (D-E) Whole-cell patch-clamp recordings from PND14-21 S1 L2/3 pyramidal neurons. Input-output firing curves (D) and rheobase and capacitance measurements (E) demonstrate reduced intrinsic excitability in Syngap1 heterozygous neurons relative to controls (Syngap1+/+ N=17 neurons, N=3 mice, Syngap1+/- N=16 neurons, N=4 mice). (F) Basal dendritic length of layer 5 pyramidal neurons in S1 at PND14. In contrast to L2/3 neurons, Syngap1 deficiency is associated with increased basal dendritic length in LS neurons, indicating lamina-specific and population-dependent structural effects (Syngap1+/+ N=46 neurons, N=S mice, Syngap1+/-N=44 neurons, N=S mice). (G) Sparse, cell-autonomous deletion of Syngap1 in S1 L2/3 pyramidal neurons using AAV-Cre in Syngap1 floxed; Ai9 reporter mice. Cell-autonomous Syngap1 loss recapitulates reduced dendritic length observed in germline mutants, indicating that structural maturation effects in sensory L2/3 neurons are intrinsic (Syngap1+/+ N=22 neurons, N=4 mice, Syngap1+/fl N=23 neurons, N=4 mice). Statistical comparisons for morphological and intrinsic properties were performed between genotypes within each region. Pairwise comparisons were assessed using two-tailed independent samples t tests unless otherwise noted. Input-output curves were analyzed using mixed-effects models fit with restricted maximum likelihood estimation (REML). Data are mean± SEM. *p < 0.05; **p < 0.01; ***p < 0.001.

We next determined to what extent impaired L2/3 cellular features could be attributed directly to intrinsic *Syngap1* deficiency (e.g. cell autonomy), rather than through indirect circuit/network-level homeostatic effects. Moreover, we sought to distinguish between cell autonomy at the population level (whole cortex-restricted excitatory neuron loss using the *Emx1*-Cre driver line) and cell autonomy at the single-neuron level by utilizing sparse *Syngap1* deletion in cKO mice using a Cre-expressing virus. Throughout, we use ‘cell-autonomous’ strictly to refer to intrinsic, single-neuron effects revealed by sparse deletion, whereas *Emx1*-Cre experiments test cortex-wide excitatory neuron restriction (cell-type, multi-regional specificity without isolating single-neuron autonomy). This approach allowed us to separate intrinsic from emergent circuit effects. Sparse induction of *Syngap1* haploinsufficiency in developing conditional *Syngap1*-OFF mice revealed that L2/3 glutamatergic neurons in primary sensory cortex also exhibited stunted dendritic arborization **(Fig. 2G)**. Whether these intrinsic excitability differences reflect cell-autonomous effects of *Syngap1* deficiency was addressed in a multi-region analysis described below (Fig. S7). These findings identify plausible developmental cellular substrates contributing to widespread depression of sensory-evoked cortical activity in fully mature sub-networks in *Syngap1* mutants. However, they also highlight a clear dissociation between widespread sensory processing deficits shown here and the systems-level hyperexcitability and seizures observed in both rodents and humans with *Syngap1* deficiency [22, 23, 25]. This contrast suggests that *Syngap1* loss does not uniformly dampen cortical function, but instead differentially affects distinct functional subnetworks.

### Syngap1 haploinsufficiency amplifies state-linked cortical responses during movement transitions

The coexistence of widespread sensory hypofunction (this study) with reported behavioral hyperactivity [24, 25] led us to ask whether *Syngap1* deficiency exerts opposing effects on cortical activity associated with internal state transitions. Spontaneous movement engages a brain-wide, state-linked cortical network that integrates arousal, motor preparatory signals, and ongoing sensory input [4–6]. If *Syngap1* deficiency differentially affects cortical subnetworks, then activity within this movement-linked state network may be differentially impacted compared to isolated sensory networks. To test this, we measured widefield calcium signals during self-initiated spontaneous movements in germline *Syngap1* mice **(Fig. 3A)**. Spontaneous movement epochs were first isolated and then segregated into small, mid, and large categories based on their total motion energy **(Fig. 3B-C**; *Supplemental Video 1*). Averaging hundreds of movements per category resulted in similar movement dynamics between genotypes during the initial post-onset window used to quantify neural responses **(Fig. 3D, G, J)**. Aligning GCaMP signals to movement onset enabled visualization of cortical activity during state transitions across movement magnitudes **(Fig. 3E, H, K)**. Consistent with past reports, the amplitude of activity across this cortical subnetwork scaled with the magnitude of movement during the state transition [4]. Comparing genotypes, there was substantially more cortical activity in *Syngap1*-deficient mice during movement transitions despite comparable motion energy **(Fig. 3F,I,L)**. Because motion energy captures overt kinematics but not internal state, these larger ΔF/F transients are consistent with enhanced movement-transition-linked cortical responses beyond what is explained by kinematics alone. Enhanced activity was most pronounced along the medial and frontal axis, with additional increases in parietal sensory regions during larger movements. Thus, in contrast to sensory-evoked distributed cortical activity, *Syngap1* haploinsufficiency is associated with amplification of state-linked cortical responses.

**Figure 3.**
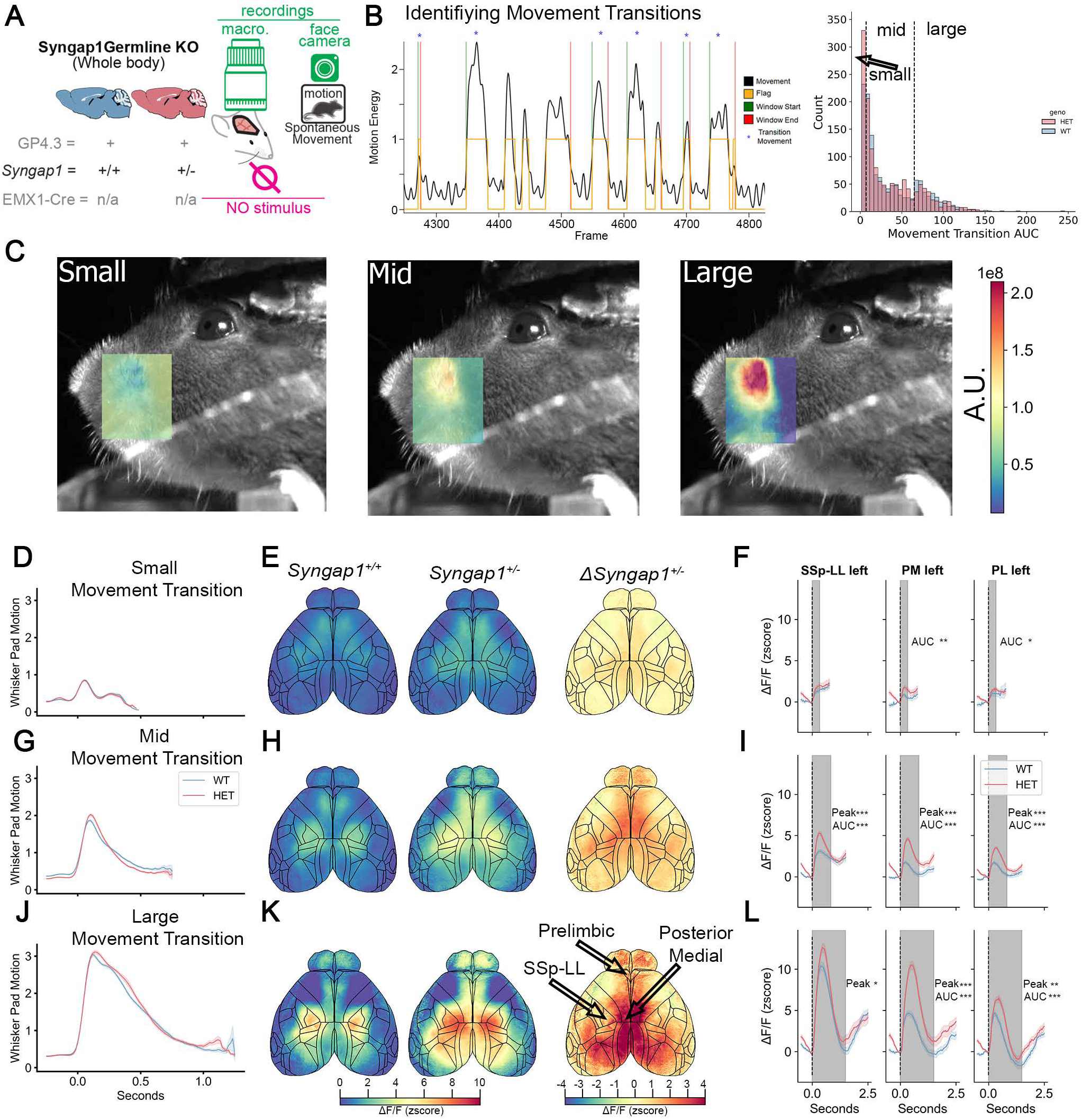
Syngap1 germline haploinsufficiency amplifies state-linked cortical responses during movement transitions. **(A)** Experimental schematic illustrating widefield GCaMP6 imaging of dorsal cortex in awake, head-fixed mice during spontaneous behavior. **(B)** Movement transitions were identified using video-derived motion energy and aligned to movement onset to isolate neural activity associated with stationary-to-movement state transitions. Only movements with clearly defined onset from quiescence were included (see Methods). Movement transitions were classified into small, medium, and large categories based on total motion energy and pooled across genotypes. **(C)** Motion energy heatmaps for control animals averaged within each movement category. Motion energy traces confirm comparable kinematics between genotypes. **(D, G, J)** Population-averaged motion energy traces for small, medium, and large movement transitions in wild-type and germline Syngap1 heterozygous mice, aligned to movement onset. (E, H, K) Widefield LlF/F activity maps aligned to movement onset for each movement category. Movement transitions engaged a distributed pattern of cortical activation spanning sensory, motor, and medial associational territories in both genotypes. (F, I, **L)** Quantification of movement-aligned cortical LlF/F responses across selected cortical regions (left lower limb somatosensory cortex: SSp-LL left, left posterior medial cortex: left PM and left prelimbic cortex: PL left). Syngap1 heterozygous mice exhibit enhanced movement-transition-linked cortical responses relative to wild-type controls despite comparable motion energy. Significance was assessed for AUC and peak amplitude (Peak) between genotypes for each cortical region and transition size within the region indicated by the shaded regions (small: 0-333ms, mid: 0-833ms, large: 0-1.Ss). Syngap1+/+ N=7 mice, small transitions N=260, mid transitions N=736, large transitions N=321. Syngap1+/- N=7 mice, small transitions N=359, mid transitions N=766, large transitions N=288. Significance was assessed using a Mann-Whitney U test with a Bonferroni correction for multiple comparisons. Data are mean ± SEM. *, p < 0.05; **, p < 0.01, *** p < 0.001.

Together with the sensory hypofunction described above, these findings establish a bidirectional mesoscale signature in *Syngap1* haploinsufficient mice: reduced sensory-evoked gain during quiescence and amplified movement/state-linked activity during transitions. We next asked whether this state-linked amplification arises from *Syngap1* loss in cortical excitatory neurons or requires broader developmental perturbation.

### Cortex-restricted Syngap1 loss reveals scope-dependent reorganization of state-linked cortical networks

Germline *Syngap1* haploinsufficiency perturbs multiple neuronal populations across cortex and subcortex, whereas *Emx1*-Cre-restricted haploinsufficiency selectively disrupts cortical excitatory neurons. Given the widespread state-linked amplification observed in germline mutants, we next tested whether restricting *Syngap1* loss to developing cortical excitatory neurons reproduces this phenotype. Using the same analytical framework applied to germline mice (Fig. 3A), we quantified cortical activity during spontaneous movement transitions in *Emx1*-Cre *Syngap1* heterozygotes. Movement kinematics were comparable across genotypes **(Fig. 4A–C)**, enabling direct comparison of neural responses. In germline mutants, movement transitions produced widespread amplification of cortical activity, most prominently along the medial and frontal axis, with comparatively smaller changes in trunk and limb somatosensory regions **(Fig. 3F,I,L; Fig. 4D)**. In contrast, *Emx1*-restricted mutants showed a distinct pattern of activity during movement transitions **(Fig. 4E–J)**. Trunk and limb somatosensory regions exhibited reduced activity that closely tracked movement dynamics, whereas medial frontal regions remained hyperactive during movement and showed a blunted return to baseline at movement offset. This resulted in persistent, spatially restricted hyperactivity extending into motor cortex. Similar patterns were observed in temporally averaged cortical maps spanning >1 s after movement onset, which showed largely preserved or modestly reduced responses across most regions, except for medial frontal areas that remained elevated **(Fig. 4E–K)**. Thus, cortical excitatory neuron loss alone is insufficient to reproduce the global state-linked amplification observed in germline mutants.

**Figure 4.**
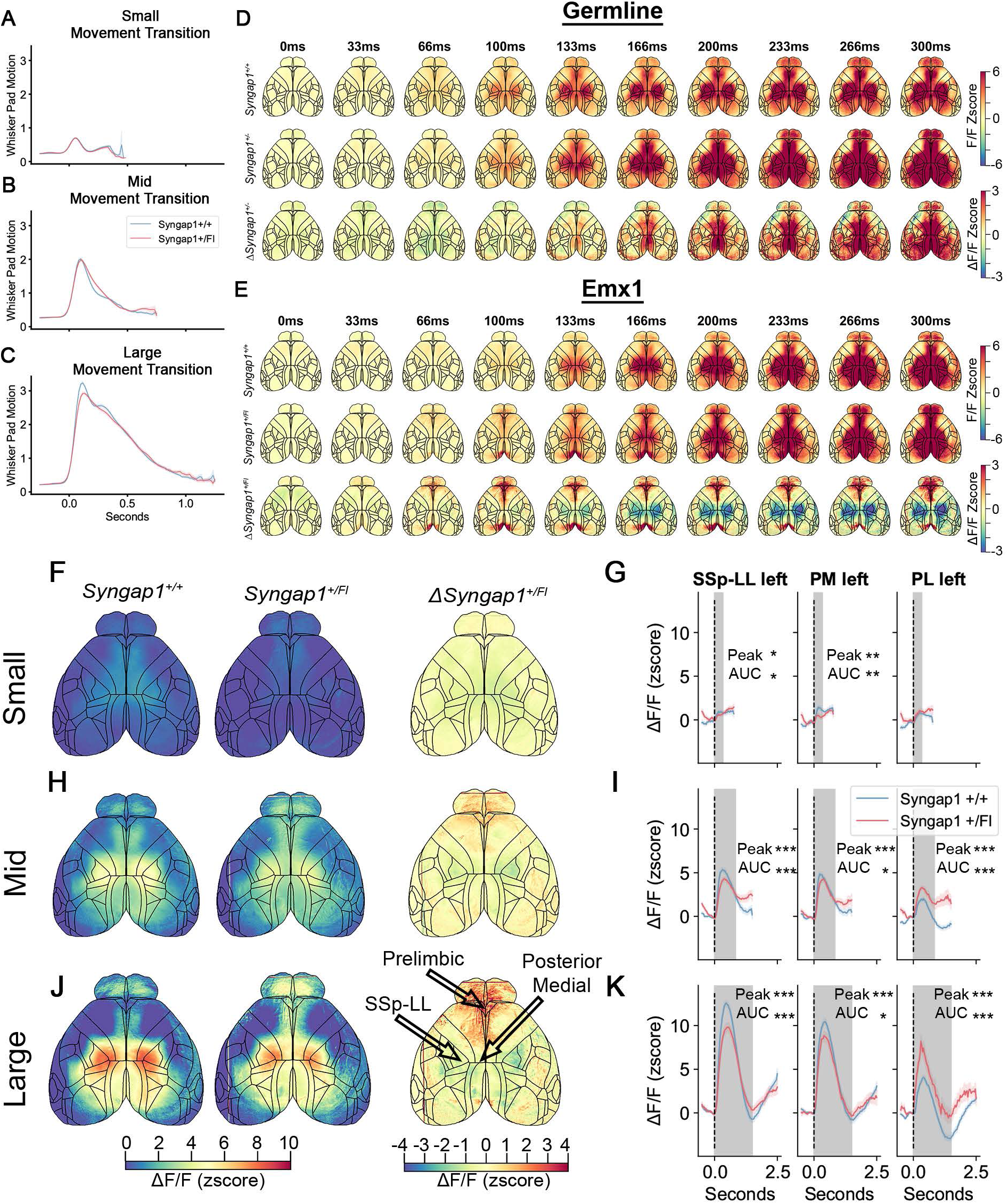
Germline and cortex-restricted Syngap1 deficiency produce distinct state-linked network reorganization. **(A-C)** Small (A), mid (B) and large (C) spontaneous movement transition motion energy traces used to identify spontaneous movement epochs and align neural activity to movement onset. (D-E) Time series of germline (D) and Emx1-restricted mutants (E) with wildtype (top), Syngap1 mutant (middle) and difference (bottom) maps showing a single frame every 33ms from 0-300ms post transition onset. **(F-G)** Time-compressed widefield GCaMP6 activity across dorsal cortex aligned to movement onset (F). Movement-aligned b..F/F signals from three representative cortical regions during small movements (G). **(H-1)** Medium-amplitude spontaneous movements, shown as in (F-G). Time-compressed widefield cortical activity aligned to movement onset (H), and movement-aligned b..F/F signals from the same cortical regions (I). **(J-K)** Large-amplitude spontaneous movements, shown as in (F-G). Time-compressed widefield cortical activity aligned to movement onset (J), and movement-aligned b..F/F signals from the same cortical regions (K). Across movement magnitudes, movement kinematics were comparable across genotypes, enabling direct comparison of neural activity associated with movement-related state transitions. Together, these analyses show that Syngap1 loss reveals dissociable patterns of state-linked co-activation that depend on the cellular scope of gene disruption. Germline Syngap1 haploinsufficiency produces a mixture of increased and decreased state-linked co-activation across dorsal cortex, whereas restricting Syngap1 loss to cortical excitatory neurons yields a distinct pattern dominated by reduced inter-regional co-activation. Significance was assessed for **AUC** and peak amplitude (Peak) between genotypes for each cortical region and transition size within the region indicated by the shaded regions (small: 0-333ms, mid: 0-833ms, large: 0-1.Ss). Syngap1+/+ N=8 mice, small transitions N=417, mid transitions N=651, large transitions N=373. Syngap1+/fl N=8 mice, small transitions N=413, mid transitions N=577, large transitions N=275. Significance was assessed using a Mann-Whitney U test with a Bonferroni correction for multiple comparisons. Data are mean± SEM. *, p < 0.05; **, p < 0.01, *** p < 0.001.

We next examined how *Syngap1* deficiency alters patterns of cortical co-activation during movement-related state transitions. Region-seeded analyses revealed distinct patterns across models **(Fig. S6A–B)**. Germline mutants exhibited a mixture of increased and decreased co-activation across dorsal cortex relative to controls **(Fig. S6A)**, consistent with heterogeneous changes in distributed networks. In contrast, *Emx1*-restricted mutants showed predominantly reduced co-activation across seeds **(Fig. S6B)**, indicating weakened interregional coupling rather than amplification.

Together, these findings demonstrate that the structure of state-linked network reorganization depends on the cellular scope of gene disruption. Global *Syngap1* haploinsufficiency produces widespread amplification and redistribution of network activity, whereas cortex-restricted loss yields a more constrained reduction in interregional coordination. These data show that *Syngap1* loss generates dissociable sensory-evoked and state-linked network phenotypes, and that the state-linked component is not explained by cortical excitatory neuron dysfunction alone. This scope-dependent dissociation indicates that distinct neuronal populations contribute differently to the observed network mosaic. We therefore asked whether region-dependent, opposite-direction shifts in developmental maturation of these populations could account for these systems-level signatures.

### Region-dependent structural and physiological maturation states are compressed in Syngap1 mutants

We next asked whether movement-linked hyperfunction in *Syngap1*-deficient mice is associated with altered maturation of L2/3 pyramidal neurons in regions engaged during state-linked activity. Prior work demonstrated accelerated dendritic maturation in frontal cortex during early postnatal development in germline *Syngap1* mice [27]. To determine whether this phenotype extends to other nodes within the state-linked network, we examined dendritic morphology of L2/3 pyramidal neurons in retrosplenial cortex (RSG/RSV) at PND14 **(Fig. 5A–B)**. Similar to frontal cortex, L2/3 neurons in RSP exhibited accelerated dendritic maturation relative to wild-type littermates. In contrast, primary sensory cortex at the same developmental stage exhibited reduced dendritic complexity (Fig. 2B,C,G). Thus, during a shared postnatal window, *Syngap1* haploinsufficiency produces opposite-direction shifts in dendritic maturation across distinct cortical regions.

**Figure 5.**
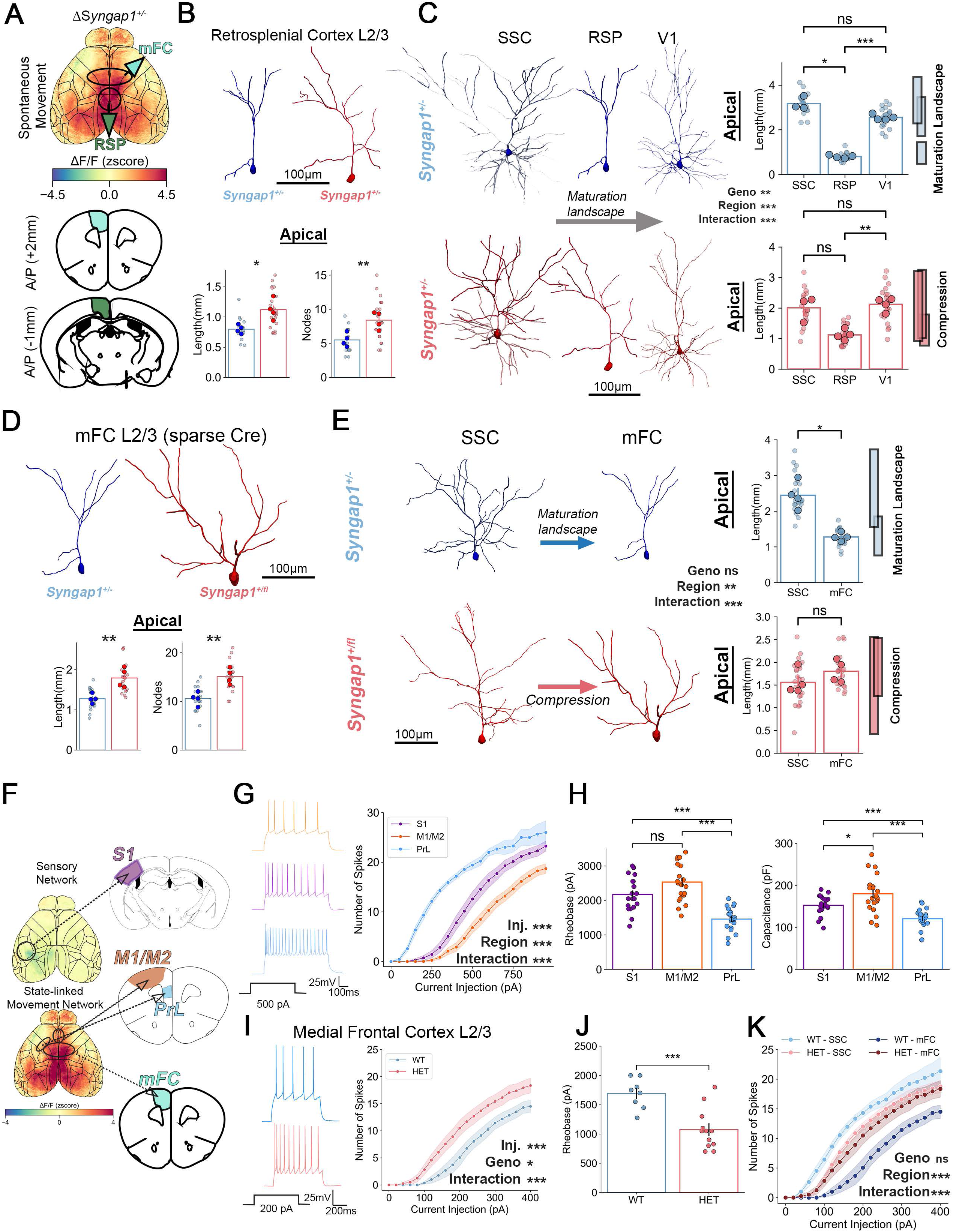
Syngap1 deficiency produces region-dependent, opposing effects on dendritic maturation and intrinsic excitability, reducing separability across L2/3 IT populations. **(A)** Schematic illustrating cortical regions selected for cellular analyses based on their engagement in state-linked movement activity identified by widefield imaging, including retrosplenial cortex (RSP) and medial frontal cortex (mFC). Accelerated dendritic maturation in frontal cortex was previously described (Aceti et al., 2015). (B) Representative dendritic reconstructions of L2/3 pyramidal neurons in retrosplenial cortex (RSG/RSV) from PND14 germline Syngap1 heterozygous mice along with quantification of apical dendrite length and node number (Syngap1+/+ N=16 neurons, N=4 mice, Syngap1+/- N=23 neurons, N=4 mice). (C) Comparative analysis of dendritic morphology in wild-type and Syngap1 heterozygous L2/3 pyramidal neurons from primary somatosensory cortex (S1), retrosplenial cortex (RSP) and primary visual cortex (V1) at matched postnatal stages (PND14). Data are replotted from Fig. 2 (S1, V1) and this figure (RSP) for direct comparison. In wild-type cortex, S1 and V1 neurons exhibit greater total dendritic length than RSP neurons, occupying separable positions within a morphological maturation landscape (highlighted by the distribution boxes on the right). In Syngap1 heterozygous mice, shortened S1 and V1 dendrites and lengthened RSP dendrites reduce separability between populations (highlighted by the compressed distributions on the right). All reconstructions were acquired and analyzed using identical imaging and tracing pipelines by experimenters blinded to genotype. (D) Sparse, cell-autonomous deletion of Syngap1 in medial-frontal L2/3 pyramidal neurons increases dendritic length at PND14, consistent with accelerated maturation in associational territories (Syngap1+/+ N=20 neurons, N=4 mice, Syngap1+/fl N=21 neurons, N=4 mice). (E) Comparative analysis of dendritic morphology in wild-type and Syngap1 heterozygous L2/3 pyramidal neurons from S1 and medial frontal cortex (mFC) at matched developmental stages. In wild-type cortex, S1 and mFC neurons are morphologically separable; sparse Syngap1 disruption reduces this separability (highlighted by separated and compressed distributions on the right). (F) Cortical territories sampled for intrinsic excitability measurements, including S1, lateral motor cortex (M1/M2), and prelimbic cortex (Prl). (G) In wild-type mice (PND14-21), L2/3 pyramidal neurons from S1, M1/M2, and Prl exhibit distinct input-output firing relationships, indicating regional differentiation in intrinsic excitability (N=22 neurons per region, N=66 neurons total). (H) Passive membrane properties from the same neurons reveal corresponding regional differences in intrinsic electrical characteristics. **(1-K)** Whole-cell patch-clamp recordings from L2/3 pyramidal neurons in medial frontal cortex (mFC) of PND14-21 germline Syngap1 mice. Representative firing responses (I) and rheobase measurements (J) demonstrate increased intrinsic excitability in mFC neurons of Syngap1 heterozygotes (Syngap1+/+ N=8 neurons, N=4 mice, Syngap1+/- N=12 neurons, N=3 mice). (K) Direct comparison of intrinsic excitability between sensory (S1) and medial frontal (mFC) L2/3 pyramidal neurons in wild-type and Syngap1 heterozygous mice. S1 data were originally plotted in Figure 2B and identical approaches were applied to both territories. In wild-type cortex, S1 and mFC populations exhibit separable firing relationships; in Syngap1 heterozygotes, these relationships converge, reflecting reduced separability of intrinsic excitability across cortical territories. Statistics for within region pairwise comparisons were assessed using two-tailed independent samples t tests. Statistics for interregional comparisons were done using ordinary two-way or three way-ANOVAs with follow up Bonferroni corrected multiple comparisons except for the interregional comparison of excitability in wild-type neurons which used a Brown-Forsyth and Welch two-way ANOVA with follow up Dunnett’s T3 multiple comparisons. Data are mean± SEM. Ns, p > 0.05, *, p < 0.05; **, p < 0.01, *** p < 0.001.

To determine whether these opposing effects reflect disruption of an underlying organizational structure in wild-type cortex, we performed a direct comparative analysis of dendritic morphology across sensory and associational regions at matched developmental stages. All reconstructions were acquired and analyzed using identical pipelines by experimenters blinded to genotype. We operationalize the “maturation landscape” as the relative ordering and separability of L2/3 IT morphological metrics across sampled cortical populations at a defined developmental time point. In wild-type mice, L2/3 pyramidal neurons in S1 exhibited substantially greater total dendritic length than neurons in retrosplenial cortex, with minimal overlap between populations **(Fig. 5C)**, indicating that cortical regions occupy distinct maturation states. In *Syngap1* heterozygous mice, this separability was reduced: shortened dendrites in sensory cortex and lengthened dendrites in retrosplenial cortex resulted in substantial overlap between populations. Thus, *Syngap1* deficiency diminishes the normal relative differentiation of dendritic maturation states across spatially separated cortical regions.

We next asked whether these region-dependent morphological effects are cell-autonomous. Sparse deletion of *Syngap1* in medial-frontal L2/3 pyramidal neurons was sufficient to increase dendritic length **(Fig. 5D)**, consistent with accelerated maturation in associational regions. Together with sparse deletion in sensory cortex (Fig. 2G), these findings demonstrate opposite-direction, cell-autonomous regulation of dendritic growth across cortical regions. Notably, sparse *Syngap1* disruption reduced morphological separability between sensory and medial-frontal populations **(Fig. 5E)**, recapitulating the convergence observed in germline mutants. These data indicate that *Syngap1* gene dosage directly influences the relative positioning of interacting cortical populations within developmental state space.

Having established region-dependent differences in dendritic maturation, we next asked whether intrinsic excitability exhibits a similarly structured maturation landscape. We first mapped the cortical regions sampled for intrinsic excitability measurements **(Fig. 5F)**. In wild-type mice (PND14–21), L2/3 neurons from S1, lateral motor cortex, and prelimbic cortex exhibited clearly separable input-output firing relationships and membrane excitability, with corresponding differences in passive membrane properties **(Fig. 5G-H)**. These data indicate that intrinsic excitability is also regionally differentiated during development.

We then asked whether germline *Syngap1* deficiency alters the relative relationships among developing populations. Medial frontal neurons in PND14-21 germline *Syngap1* heterozygous mice were hyperexcitable relative to wild-type controls **(Fig. 5I-J)**. To assess effects of *Syngap1* deficiency on relative separation of excitability across regions, we again performed a comparative analysis, where data from Fig. 2D (S1) and Fig. 4I (mFC) were plotted together. All recordings were performed using identical acquisition and analysis pipelines, with experimenters blinded to genotype. In wild-type mice, we observed that L2/3 pyramidal neurons from S1 and medial-frontal regions exhibited clearly separable input-output curves **(Fig. 5K),** where S1 neurons exhibited greater excitability compared to medial frontal neurons, a finding consistent with our prior experiment *(Fig. 4G)*. However, in *Syngap1* heterozygous mice, this separation was lost, with firing rate curves from the two populations converging and substantially overlapping **(Fig. 5K),** reflecting loss of regional differentiation in intrinsic excitability.

To determine whether intrinsic excitability differences are cell-autonomous, we performed sparse *Syngap1* deletion across S1 and frontal regions **(Fig. S7A)**. In contrast to dendritic morphogenesis, sparse deletion did not alter intrinsic excitability in any cortical region examined **(Fig. S7B)**. Regional excitability differences observed in germline wild-type mice were replicated in Cre-negative neurons from *Syngap1* flox mice **(Fig. S7C)**, supporting the stability of region-specific excitability landscapes. These findings indicate that intrinsic excitability differences across L2/3 populations emerge from circuit-level context rather than single-neuron intrinsic programs alone.

Together, these findings demonstrate that *Syngap1* haploinsufficiency reduces relative differentiation across multiple developmental substrates in L2/3 IT populations. Dendritic structure is regulated cell-autonomously in opposite directions across cortical regions, whereas intrinsic excitability landscapes, though similarly convergent in mutants, depend on circuit context. Reduced *Syngap1* expression therefore diminishes the normal separation of maturation states across spatially distributed L2/3 populations. These opposing developmental shifts are directionally aligned with the sensory-evoked hypofunction and state-linked hyperfunction observed at the mesoscale, consistent with a developmental biasing mechanism underlying the emergence of distributed network imbalance.

### ERK signaling is differentially engaged across cortical regions following Syngap1 loss

Having established that *Syngap1* haploinsufficiency compresses normally distinct dendritic and excitability landscapes across L2/3 IT populations, and that intrinsic excitability differences are not cell-autonomous, we next asked whether these altered maturation states are associated with differential engagement of conserved intracellular signaling pathways. SynGAP is a Ras-MAPK pathway regulator [35], and ERK signaling is well positioned to influence intrinsic excitability through modulation of membrane conductances and downstream transcriptional programs [36, 37]. We therefore tested whether *Syngap1* deficiency alters sensitivity to ERK pathway inhibition in a region-specific manner, which could reveal a molecular correlate consistent with the compressed maturation landscape observed across cortical L2/3 IT populations. To address this, we prepared organotypic slice cultures from wild-type and *Syngap1* heterozygous mice and performed whole-cell patch-clamp recordings from L2/3 pyramidal neurons in primary somatosensory cortex (S1) and medial frontal cortex (mFC; **Fig. 6A**; *see also Fig. 5A*). Organotypic cultures from cortex preserve local microcircuit architecture while permitting multi-day pharmacological manipulation, allowing ERK signaling to be interrogated within an intact developmental circuit context in which excitability landscapes emerge. Slices were from neonates cultured for at least a week in the presence of vehicle (DMSO) or an established MEK inhibitor (PD98059; 10μM) to persistently reduce ERK1/2 signaling **(Fig. 6A)**. Across 152 independent whole cell recordings, several findings emerged. Initially, we confirmed that region-specific intrinsic excitability phenotypes observed in acute slices were reproduced in cultured preparations. Consistent with earlier findings, L2/3 pyramidal neurons in S1 were hypoexcitable in *Syngap1* mutants relative to wild-type controls **(Fig. 6B-C)**, whereas L2/3 neurons in mFC were hyperexcitable **(Fig. 6F-G)**, confirming preservation of opposing excitability states in vitro. In wild-type neurons, ERK inhibition revealed region-specific baseline roles. In S1, ERK inhibition significantly reduced wild-type intrinsic excitability, as evidenced by reduced spike output and increased rheobase **(Fig. 6B-C)**, indicating that ERK signaling in development contributes to excitability in this population. In contrast, Mek inhibition had no detectable effect on current-spike relationships or rheobase in wild-type mFC neurons **(Fig. 6F-G)**, suggesting that ERK signaling does not tonically regulate excitability in this region at this developmental stage. Strikingly, *Syngap1* deficiency revealed genotype-dependent inversion of ERK-sensitive excitability across regions. In S1 neurons from *Syngap1* heterozygous mice, ERK inhibition increased intrinsic excitability, shifting responses toward wild-type levels **(Fig. 6B-C)**. Statistical analysis revealed a significant genotype × treatment interaction for rheobase in S1 neurons, with post hoc comparisons demonstrating normalization of mutant responses following ERK inhibition **(Fig. 6C)**. A similar bidirectional interaction was observed for resting membrane potential **(Fig. 6D)**, but not membrane resistance **(Fig. 6E)**, indicating that ERK-dependent control of a subset of intrinsic membrane properties is altered in a genotype-dependent manner in sensory cortex. In mFC neurons from *Syngap1* heterozygous mice, MEK inhibition significantly reduced intrinsic hyper-excitability leading to firing responses no different than wild-type levels **(Fig. 6F-G)**. Consistent with this selective effect, ERK inhibition significantly altered membrane properties, including resting membrane potential and membrane resistance, only in mutant mFC neurons and in a direction consistent with reduced excitability **(Fig. 6H-I)**. Thus, *Syngap1* loss alters the functional relationship between ERK pathway activity and intrinsic excitability in a cell-type/region x genotype dependent manner.

**Figure 6.**
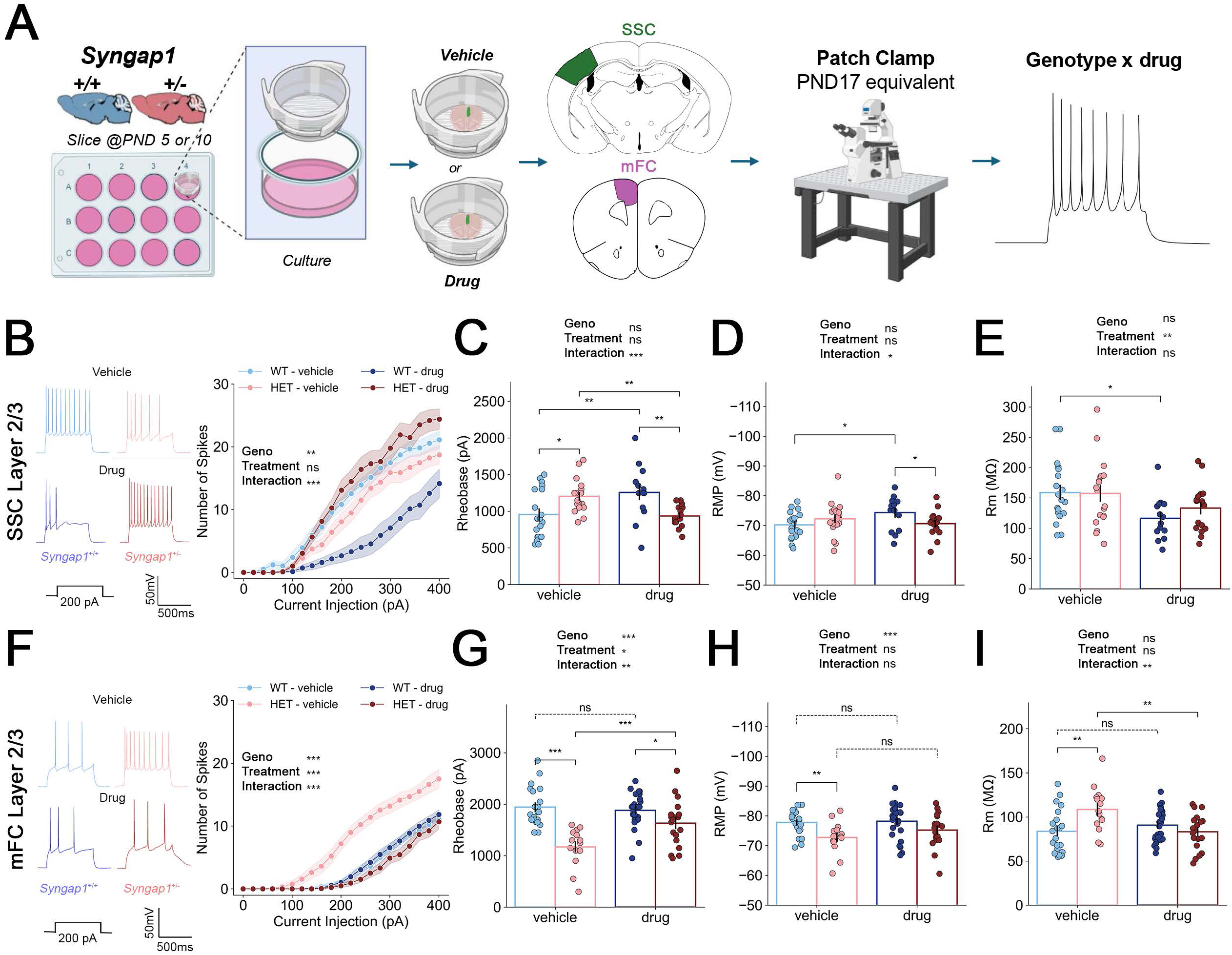
Syngap1 deficiency reveals genotype- and region-dependent inversion of ERK-sensitive excitability across L2/3 IT neurons. **(A)** Experimental design. Organotypic slice cultures were prepared from **PND 5** (medial frontal cortex, mFC) or **PND** 10 (primary somatosensory cortex, S1) wild-type and Syngap1 heterozygous mice and maintained until PND17 in the presence of vehicle or the MEK inhibitor PD98059 (10 µM), which reduces ERK1/2 pathway activity. Whole-cell recordings were obtained from L2/3 pyramidal neurons in S1 and mFC. (B-E) ERK inhibition in S1 neurons. (B) Representative current-spike input-output curves from S1 L2/3 pyramidal neurons across genotype and treatment conditions and quantification of spike output. (C) Rheobase measurements. (D) Resting membrane potential (RMP). (E) Membrane resistance. In WT S1 neurons, ERK inhibition reduced intrinsic excitability (decreased spike output, increased rheobase). In contrast, in Syngap1 HET S1 neurons, ERK inhibition increased intrinsic excitability and restored responses toward WT levels. Two-way ANOVA revealed a significant genotype _x_ treatment interaction for rheobase (see Methods and Statistics Table). (F-G) ERK inhibition in mFC neurons. (F) Representative input-output curves from mFC L2/3 pyramidal neurons and quantification of spike output. (G) Rheobase measurements. ERK inhibition had no detectable effect on intrinsic excitability in WT mFC neurons. In contrast, ERK inhibition significantly reduced intrinsic excitability in Syngap1 HET mFC neurons, normalizing firing responses toward WT levels. Two-way ANOVA revealed a significant genotype _x_ treatment interaction (see statistics). (H-I) Membrane property analyses. (H) Resting membrane potential (RMP). (I) Membrane resistance. In S1, ERK inhibition produced genotype-dependent shifts in RMP consistent with bidirectional regulation of intrinsic excitability. In mFC, ERK inhibition selectively altered membrane properties in Syngap1 HET neurons but not WT neurons. Together, these data demonstrate that ERK signaling exerts region-specific control over intrinsic excitability in WT cortex and that Syngap1 haploinsufficiency reprograms this control in opposite directions across cortical territories. The same conserved signaling pathway is associated with divergent excitability outcomes depending on cortical region and genotype, providing a molecular correlate consistent with compression of normally distinct excitability landscapes across L2/3 IT populations (see Fig. 4). Data are mean± SEM. Ns, p > 0.05, *, p < 0.05; **, p < 0.01, *** p < 0.001. Statistical tests and exact n values are provided in Methods.

Together, these findings demonstrate that *Syngap1* deficiency does not uniformly elevate or suppress intrinsic excitability through a single signaling mechanism. Instead, *Syngap1* loss reconfigures ERK-dependent control in a region-specific manner, such that the same conserved signaling pathway supports excitability in sensory cortex under baseline conditions, yet becomes aberrantly engaged to drive hyperexcitability in medial frontal cortex following gene disruption. This bidirectional signaling logic parallels the compression of normally distinct excitability landscapes across L2/3 IT populations (Fig. 5). Moreover, it provides a molecular example consistent with how a developmental regulator can differentially engage conserved intracellular pathways to produce opposing circuit-integration phenotypes across molecularly distinct L2/3 IT neurons residing in different cortical regions that functionally interact during sensation and action.

### Altered sensory–state integration biases behavioral transitions

The preceding analyses demonstrate that *Syngap1* haploinsufficiency produces opposing mesoscale functional network states, and that the direction of state-linked amplification depends on the cellular scope of gene disruption. If these network imbalances reflect altered integration rules between sensory and state-linked subnetworks, then they should bias how sensory events are translated into movement transitions. We therefore asked whether the inversion of state-linked network activity observed in germline versus *Emx1*-restricted mutants is reflected in the structure and timing of spontaneous and sensory-evoked behavior. Using Facemap [6], we again analyzed spontaneous movement bouts from animals that were extensively habituated to head fixation **(Fig. 7A)**. Because behavioral structure was the focus of this analysis, instead of identifying clear state transitions as in earlier analyses (e.g. stationary>move), all movement bouts were flagged and averaged across genotypes from both strains with much less stringent inter-movement time-outs. This analysis identified substantially more movements than in prior analyses. A *k*-means clustering analysis identified three clear movement categories, which were given the same small, mid, and large designations. “Small” movements resembled “twitches”, which were defined by abrupt, low-amplitude motion energy traces **(Fig. 7B)**. In contrast, “large” movements were much longer in duration and amplitude, and they co-occurred with whole body repositioning **(Fig. 7B)**. “Medium” movements were intermediate, and they often coincided with whisking/sniffing **(Fig. 7B)**. Averaging all movements within each category revealed relative similar motion energy dynamics between genotypes across each type of movement **(Fig. 7B)**. However, while the overall movement dynamics were highly similar between genotypes, there were qualitatively subtle deviations between genotypes at various points in the motion energy curves, either in amplitude or duration.

**Figure 7.**
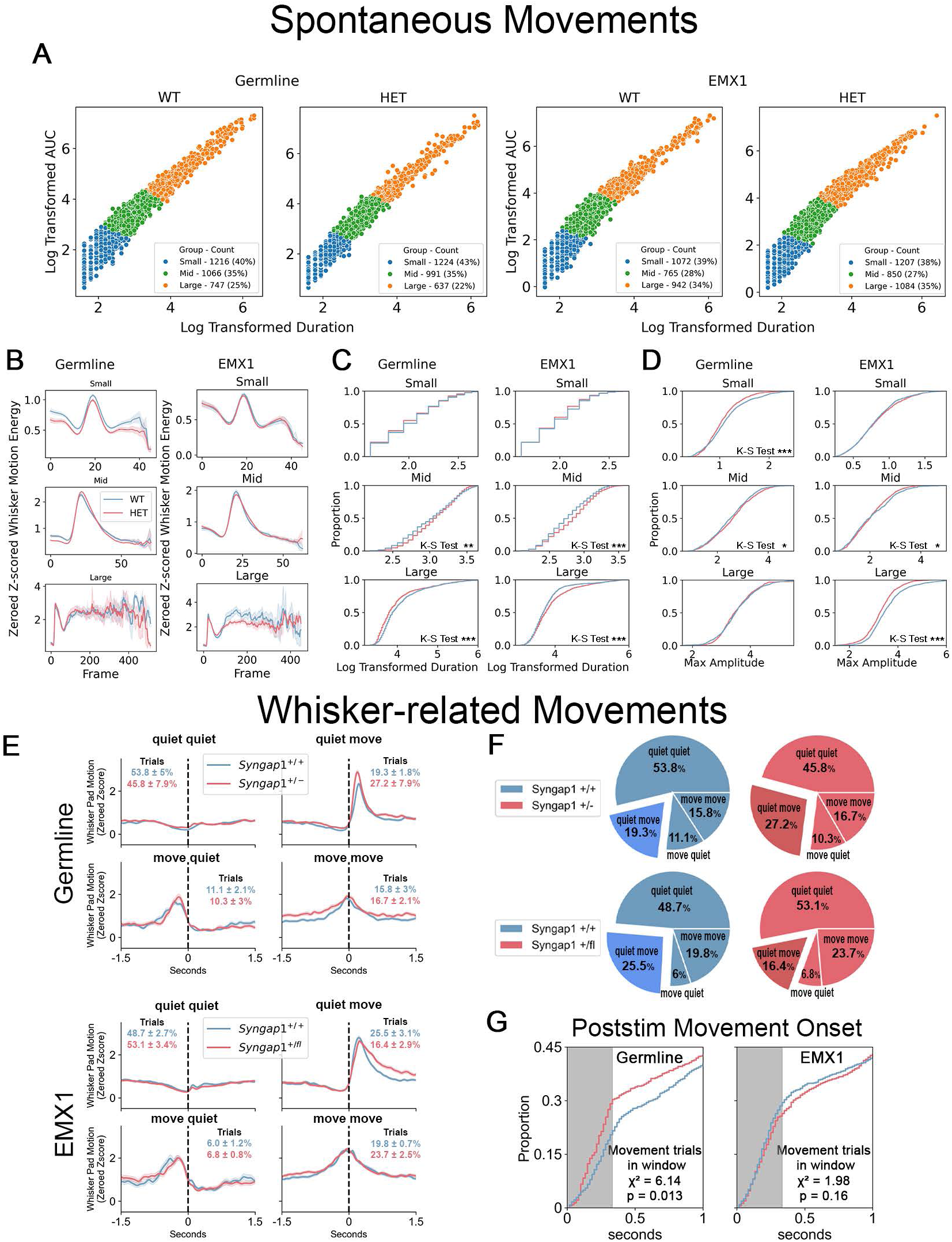
Behavioral differences in movement structure and sensory-evoked transitions align with opposing cortical activity regimes in Syngap1 mutants. **(A-D)** Classification of all spontaneous movements based on motion energy from both germline and Emx1-Syngap1 deficient models. K-means clustering (see methods) of all movements revealed three distinct movement types: small (blue), mid (green), and orange (large) movement **(A).** Representative motion energy traces illustrate the three movement categories identified by clustering. small, medium, and large movements **(B).** These categories were qualitatively consistent with the small, mid, and large movement bins defined in Figure 3 and were similar across genotypes and strains. Small movements correspond to brief, low-amplitude motion ("twitches"), medium movements often coincide with whisking or sniffing, and large movements are longer in duration and amplitude and frequently involve whole-body repositioning. Cumulative probability distributions of movement duration and amplitude for each movement category (C-D). Germline Syngap1 heterozygous mice exhibit a higher proportion of short-duration, high-amplitude movements compared to wild-type controls, whereas Emx1-Cre Syngap1 heterozygous mice have a lower proportion of high-amplitude movements (Germline: Syngap1+/+ N=7 mice, Syngap1+/- N=7 mice; Emx1-Cre: Syngap1+/+ N=8 mice, Syngap1+/fl N=8 mice). (E-F) Analysis of movement responses during whisker-evoked trials. Sensory trials were categorized based on movement occurrence before and after whisker stimulation (quiet-quiet, quiet-move, move-quiet, move-move). Germline and EMX1-Syngap1 heterozygous mice show distinct sensory-evoked movement kinematic profiles (E) and movement probabilities (F) relative to controls during quiet-move trials. (G) Cumulative probability of time to first movement following whisker stimulation in each genotype from both models. Germline Syngap1 heterozygous mice exhibit earlier movement onset relative to wild-type littermates, whereas Emx1-Cre Syngap1 heterozygous mice show delayed movement responses. Together, these analyses indicate that the opposing cortical activity regimes observed at the mesoscale are associated with distinct alterations in spontaneous movement structure and sensory-evoked movement timing (Germline: Syngap1+/+ N=1399 trials, N=7 mice, Syngap1+/- N=1008 trials, N=S mice; Emx1-Cre: Syngap1+/+ N=1200 trials, N=8 mice, Syngap1+/fl N=1200 trials, N=8 mice). Data are mean± SEM. *, p < 0.05; **, p < 0.01, *** p < 0.001.

This finding suggested that subsets of movements within each category may be uniquely impacted by *Syngap1* deficiency, including both the germline and the *Emx1*-restricted models. To test this, we defined each movement from each animal by its duration and amplitude. All movements for each category in each genotype were then plotted on cumulative probability curves **(Fig. 7C-D)**. This analysis revealed how specific types of movements were impacted by *Syngap1* deficiency. Germline mutants had a significantly higher proportion of small movements that were lower amplitude, while they also exhibited large movements with shorter duration compared to littermate controls. In contrast, while *Emx1*-restricted *Syngap1* mutants also exhibited different movements compared to their controls, the directionality of the effects tended to be in the opposite direction compared to germline mutants. For example, large movements in *Emx1* mutants were significantly lower in amplitude but longer in duration compared to their littermate controls.

We next examined movement properties during whisker-evoked trials **(Fig. 7E-F)**, which also revealed an inversion of movement characteristics between the two *Syngap1* models. First, we assessed the probability that a movement occurred in response to a whisker stimulation. We did this by segregating all head-fixed whisker-evoked sensory trials into four categories based on movements before and/or after the whisker stimulation (e.g., quiet-quiet, quiet-move, move-quiet, move-move; **Fig. 7E,F-G)**. Because mice are sensitive to whisker deflections, we reasoned that quiet-move trials reflected sensory-driven movement transitions. Germline *Syngap1* mutants had significantly more *“quiet-move”* trials compared to wildtype littermates [Fishers exact test: WT (N=7 mice): 270 vs. 1130 (total); HET (N=5 mice): 274 vs. 734 (total), Stat: 0.64, p<10-e5]. In contrast, EMX1-Hets had significancy fewer “*quite-move*” trials (Fishers exact test: WT (N=8 mice): 306 vs. 894 (total); Emx1-HET (N=8 mice): 197 vs. 1003(total), Stat: 1.74, p<10-e8]). Second, for all whisker stimulus trials, we time-stamped the first movement following stimulus onset and plotted the cumulative probability of post-stimulus movements **(Fig. 7G)**. Treating stimulus onset as time = 0 allowed us to quantify the temporal bias toward movement initiation during sensory processing. Germline *Syngap1* heterozygotes exhibited significantly earlier movement initiation compared to wild-type littermates, whereas *Emx1*-restricted heterozygotes showed a trend toward delayed movement relative to stimulus onset. Together, these findings indicate that opposing functional network states in germline and cortex-restricted *Syngap1* mutants are associated with directionally biased sensory-to-movement transitions, consistent with model-specific alterations in integration between sensory and state-linked subnetworks.

## DISCUSSION

Neurodevelopmental disorders are increasingly conceptualized as disorders of distributed brain systems rather than focal circuit defects [38, 39]. Here, we identify a developmental mechanism through which a single risk gene can differentially shape interacting cortical populations to organize distributed network architecture. Using *Syngap1* haploinsufficiency as a causally defined perturbation, we demonstrate coexisting, opposing cortical activity states within the same animal model: sensory-evoked responses are reduced, whereas movement/state-linked activity is amplified. These divergent mesoscale signatures are accompanied by corresponding behavioral shifts, indicating that *Syngap1* loss disrupts distributed network balance rather than uniformly altering excitability.

A central finding is that these opposing network states are associated with asymmetric, region-dependent effects of *Syngap1* on neuronal maturation during a defined postnatal window of circuit assembly. While prior work established that *Syngap1* regulates maturation within individual neuronal populations during this period (Clement et al., 2012; Clement et al., 2013; Aceti et al., 2015), the present data reveal simultaneous, opposite-direction shifts across distinct cortical regions. In primary sensory cortex, *Syngap1* promotes dendritic growth and intrinsic responsiveness in Layer 2/3 IT neurons. In contrast, in medial frontal and associational regions, *Syngap1* constrains dendritic maturation and excitability. Sparse deletion demonstrates that dendritic morphogenesis is regulated cell-autonomously in opposing directions across regions, whereas intrinsic excitability differences require circuit context and are not reproduced by single-neuron deletion. Together, these findings show that a single NDD risk gene exerts pleiotropic, cell-type-specific control over core substrates of circuit integration across spatially distributed cortical populations.

These opposing cellular phenotypes driven by *Syngap1* pleiotropy converge on a systems-level model **(Fig. 8)**. In wild-type cortex, L2/3 IT populations across sensory and associational regions occupy separable maturation states defined by dendritic architecture and intrinsic excitability. *Syngap1* haploinsufficiency reduces this separability by shifting interacting populations in opposite directions, resulting in convergence of their maturation states. This pattern is consistent with hierarchical models of cortical development in which sensory and associational regions follow partially offset maturation trajectories along a sensorimotor-association axis [15, 16, 40, 41]. However, our findings extend these models by identifying a gene-dependent mechanism that can alter the relative positioning of interacting neuronal populations within this developmental axis. Here, we use “compression” to describe reduced separability of maturation states across populations, and “reweighting” to describe the observed shift in balance between sensory-evoked and movement-linked activity, without implying directed causality. We propose that distributed cortical systems are assembled through coordinated relative maturation of interacting neuronal populations. Perturbations that shift the relative developmental positioning of these populations can bias how distributed networks are integrated. Although we do not directly link developmental cellular phenotypes to adult network states within the same animals, the opposing developmental shifts observed here are directionally aligned with later subnetwork gain imbalance, consistent with a developmental biasing mechanism rather than a one-to-one mapping. Under this framework, altered relative maturation across interacting populations provides a plausible basis for the emergence of stable mosaics of hypo- and hyper-functional cortical subnetworks.

**Figure 8.**
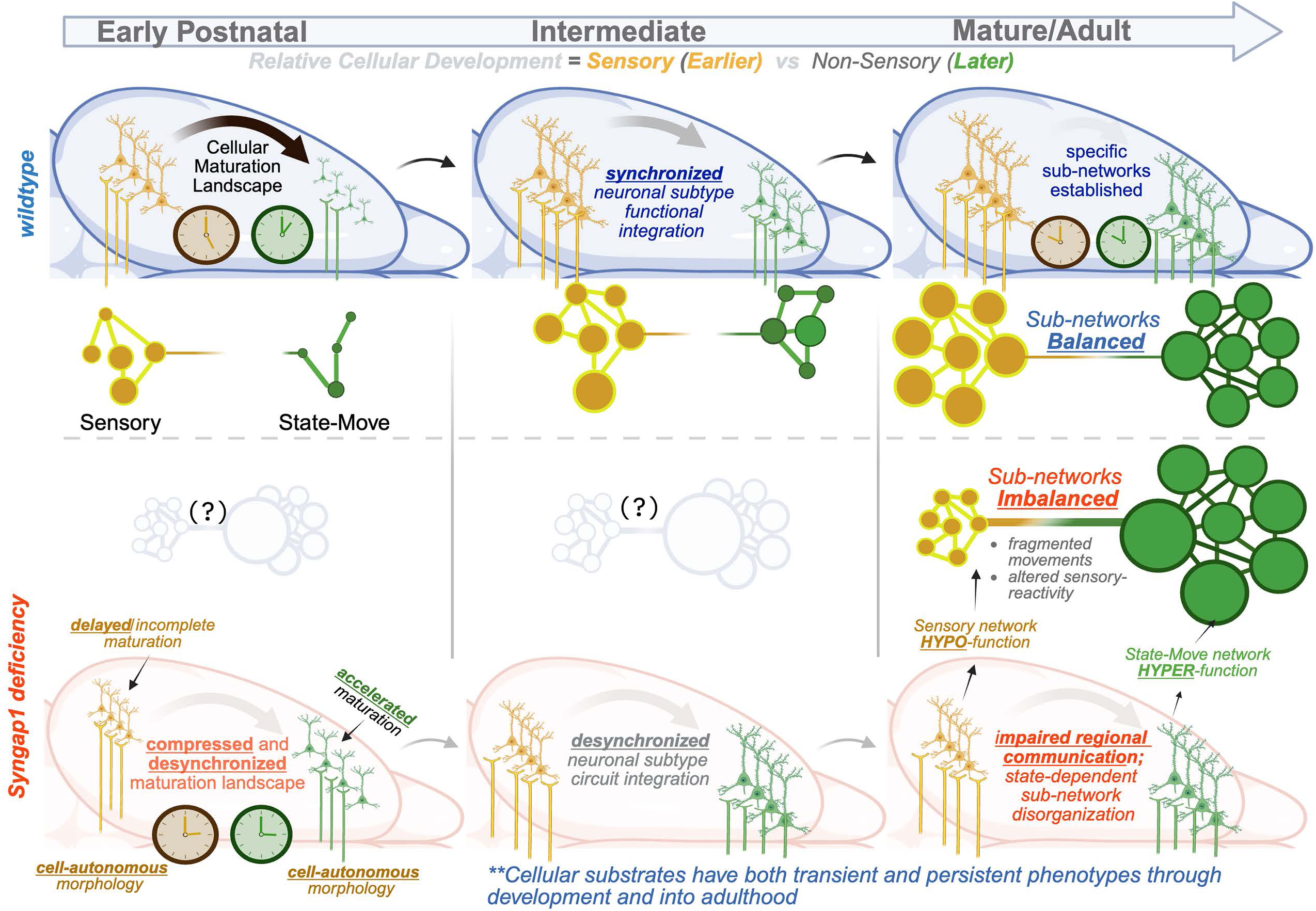
Model: Relative maturation states across cortical populations shape distributed network balance. This schematic summarizes a developmental framework in which the relative positioning of interacting neuronal populations within a maturation landscape influences the later balance of distributed cortical subnetworks. Top (wild-type): During early postnatal development, Layer 2/3 intratelencephalic (IT) neurons across sensory and associational cortex occupy separable maturation states, reflected in distinct dendritic architectures and intrinsic excitability landscapes. This relative differentiation provides a substrate through which sensory-driven and state-linked subnetworks can integrate in a balanced manner in adulthood. Bottom (Syngap1 haploinsufficiency): Loss of Syngap1 is associated with delayed maturation of sensory-coding neurons and accelerated maturation of select associational populations, reducing separability within the developmental maturation landscape. Although some cellular phenotypes may be transient, shifts in relative maturation state can bias how neuronal ensembles assemble and stabilize circuit interactions. In adulthood, this is associated with reweighted cortical subnetworks characterized by sensory hypofunction and amplified state-linked activity. We propose that distributed cortical systems are assembled through coordinated relative maturation of multiple separable neuronal populations. Gene-dependent perturbations that alter the relative positioning of these populations within developmental state space can bias the gain and integration properties of distributed networks.

Our ERK experiments provide a mechanistic illustration of region-dependent signaling during development, in which *Syngap1*–ERK pathway interactions are cell-type specific. ERK inhibition produces distinct effects across cortical regions in wild-type neurons and reveals genotype-dependent inversion of ERK-sensitive excitability in *Syngap1* mutants. In sensory cortex, ERK inhibition counteracts gene-driven hypo-excitability in *Syngap1*-deficient neurons and restores responses toward wild-type levels. In contrast, in medial frontal cortex, the same manipulation selectively normalizes intrinsic hyperexcitability in mutant L2/3 IT neurons. These effects, observed in organotypic cultures that preserve local circuit architecture, are consistent with region-specific engagement of conserved intracellular pathways rather than a single uniform signaling defect. This provides molecular evidence that a developmental regulator can differentially tune a shared signaling pathway across interacting neuronal populations.

This framework integrates prior circuit-level findings in *Syngap1*-deficient mice that appeared paradoxical when considered in isolation. In adult primary somatosensory cortex, *Syngap1* haploinsufficiency reduces feedforward synaptic drive from layer 4 to layer 2/3 pyramidal neurons, consistent with weakened sensory encoding [30]. Subsequent work demonstrated that whisker-evoked activity in layer 5 somatosensory cortex receives weakened local sensory input while simultaneously exhibiting enhanced feedback connectivity from motor cortex [26]. During active behavior, silicon probe recordings revealed coordinated increases in spiking across somatosensory cortex, motor cortex, and thalamus during whisking, whereas active touch suppressed firing in touch-sensitive neurons [26]. Rather than reflecting unrelated circuit defects, these opposing synaptic and state-dependent phenotypes are consistent with altered relative weighting of feedforward sensory and feedback state-related inputs. Within the framework summarized in **Figure 8**, such circuit-level reorganization represents a local manifestation of regionally biased maturation states that ultimately influence distributed cortical network balance. In this sense, altered relative maturation may shift the alignment and weighting of communication subspaces across regions [5, 6], biasing how sensory and state-related signals are integrated at the population level.

The use of anatomical “regions” in this study reflects an operational framework rather than strict biological boundaries. High-density electrophysiological mapping demonstrates that functional organization does not strictly follow cytoarchitectural subdivisions but instead reflects continuous gradients of activity and connectivity [17]. In this context, region-dependent effects of *Syngap1* are best interpreted as biases in maturation of functionally coupled neuronal ensembles rather than disruptions confined to discrete parcels. Retrosplenial cortex illustrates this principle: it participates in both sensory-evoked and state-linked movement networks and exhibits context-dependent activity changes following *Syngap1* loss.

Convergent evidence suggests that neurodevelopmental risk genes influence core substrates governing neuronal integration, including dendritic morphogenesis, synaptic organization, and intrinsic excitability. Single-cell transcriptomic atlases demonstrate that Layer 2/3 glutamatergic projection neurons are transcriptionally specialized across cortical areas, whereas inhibitory interneurons are comparatively conserved [42–44]. Recent *in vivo* multiomic Perturb-seq studies further show that perturbation of neurodevelopmental disorder risk genes produces highly cell-type-specific transcriptional and chromatin phenotypes across developing cortical populations, and that baseline expression alone is insufficient to predict functional impact [45]. These findings provide a molecular proof-of-concept that NDD gene function is interpreted in a context-dependent manner across neuronal populations. Our results extend this principle by showing that a single NDD gene perturbation can produce opposing developmental trajectories across L2/3 IT neurons from interacting cortical regions and that these divergent cellular outcomes map onto altered distributed network function. Together, these findings support the view that gene-dependent perturbations can differentially bias separable excitatory neuron populations, altering their relative developmental positioning and, consequently, distributed network function.

The dissociation between germline and cortex-restricted *Syngap1* mutants further constrains this model. Although both genotypes exhibit behavioral hyperactivity [24, 25], widespread amplification of state-linked cortical activity during movement transitions emerges only in germline mutants. This indicates that state-linked network hyperfunction is an emergent distributed phenotype requiring perturbation beyond cortical excitatory neurons alone. Given the known role of *Syngap1* in other cortical neuron subtypes and brain areas, inhibitory interneuron dysfunction [46] and altered striatal D2 neuron maturation [47] likely modulate how cortical maturation landscapes are expressed at the systems level. Thus, developmental coordination is shaped not only by intrinsic differentiation programs but also by interactions among excitatory, inhibitory, and subcortical populations.

Although this study focuses on *Syngap1*, the organizing framework is unlikely to be gene-specific. Many ASD-and ID-associated genes are expressed in upper-layer projection neurons during periods of rapid dendritic growth and circuit integration [19, 21]. Region- or cell-type-dependent effects on structural and physiological differentiation, even modest ones, could shift the relative positioning of interacting populations within developmental state space and bias distributed network balance. In this sense, *Syngap1* provides a mechanistically tractable example of a likely broader class of developmental regulators influencing coordination across neuronal ensembles.

Finally, this framework extends beyond autism. Large-scale genomic analyses reveal substantial shared genetic liability across diverse psychiatric conditions, including autism, schizophrenia, bipolar disorder, major depression, and attention-deficit/hyperactivity disorder [18]. Yet the developmental system mechanisms linking shared genetic risk to distributed network dysfunction remain poorly defined. This idea is also consistent with recent in vivo perturbational atlas approaches showing that NDD risk genes can produce divergent molecular effects across cortical neuronal subclasses [45]. Gene-dependent shifts in the relative developmental timing across interacting neuronal populations represent one candidate mechanism through which convergent genetic perturbations could bias large-scale network integration. Under this view, modest, region- and cell-type-specific shifts in relative differentiation across interacting populations may produce stable mosaics of network imbalance that manifest as distinct behavioral syndromes depending on timing, circuit context, and affected populations.

## Acknowledgments

The authors would like to thank Simon Musall, Ph.D. for technical discussions regarding macroscope construction, Carlos Portera-Cailliau, M.D., Ph.D. for constructive comments on an earlier version of this manuscript, and members of the Rumbaugh Lab for insightful conversations regarding data analysis and interpretations. We would also like to thank the National Institutes for Mental Health (F32MH130163 to RG; R01MH096847 and R01MH131788 to GR) and the National Institute for Neurological Disorders and Stroke (R01NS110307 to GR) for providing generous funding support for this research.

## Author Contributions

<u>Methodology:</u> RG, BGG, SDM, MA, SB, CR; <u>Validation:</u> RG, BGG, SDM, MA, SB, CR; <u>Writing - Original Draft:</u> RG, GR, BGG; <u>Writing - Review & Editing:</u> RG, BGG, SDM, MA, SB, CR, CAM, TV, GR; <u>Supervision:</u> TV, CAM, GR; <u>Conceptualization:</u> GR.

## METHODS

### Animals

All mouse experiments were carried out in accordance with the NIH Guide for the Care and Use of Laboratory Animals and approved by the Scripps/UF Scripps Biomedical Research Institutional Animal Care and Use Committee. Both male and female mice were used in all experiments. The *Syngap1* mouse lines used in this study have been previously validated [28, 48]. We used germline heterozygous *Syngap1* knock-out (*Syngap1^+/-^*, JAX#008890) and conditional *Syngap1* floxed (*Syngap1^+/fl^*, JAX#029303) mice. For germline *Syngap1* widefield calcium imaging experiments, *Syngap1^+/-^* mice were crossed to hemizygous (*Thy1*-GCamp6s)GP4.3 (*Thy1*-GCamP6s, JAX#024275) mice purchased from Jackson labs. For *Syngap1* conditional knock-out widefield calcium imaging experiments, first *Syngap1^+/fl^* mice were crossed with *Thy1*-GCamP6s to yield double transgenic animals. Next, these double transgenic *Syngap1^+/fl^*; *Thy1*-GCamP6s mice were crossed with *Emx1*-Cre mice (JAX#005628) acquired from Jackson Labs to yield the experimental *Syngap1* conditional knock-out mice. For neuronal tracing experiments, *Syngap1^+/-^* mice were bred with *td*Tomato Ai9 (Ai9, JAX#007905) to yield *Syngap1^+/-^*; Ai9^+/+^ experimental animals for germline experiments and *Syngap1^+/fl^*was crossed with Ai9 mice to yield *Syngap1^+/fl^*; Ai9^+/-^ for sparse expression experiments. For sparse expression electrophysiology Ai6-*Zs*green mice (*Zs*green, JAX#007906) were crossed with *Syngap1^+/fl^*. Experimental cohorts were generated from age-matched littermates taken from multiple litters and balanced as well as possible for genotype and sex. Experimenters remained blinded to the genotype of animals for data acquisition and analysis. All animals were maintained in a 12-hour light/dark cycle in temperature and humidity-controlled rooms.

### Widefield Surgery

The widefield surgeries were done similarly to published protocols with small adjustments [30, 49]. The dorsal surface of the skull was made optically transparent by applying cyanoacrylate glue (Zap-A-Gap medium CA+) and a custom made (MPFI machine shop) stainless steel head post was affixed to the skull of mice aged 8-10 weeks. Animals were anesthetized using isoflurane (5% induction, 1.5-2% maintenance) with a low flow vaporizer (Somno Low-Flow Vaporizor, Kent Scientific) and stabilized in a stereotaxic frame (David Kopf Instruments). Mouse body temperature was regulated by a feedback-controlled pad set at 37 ⁰C placed under the animal. Eyes were kept lubricated during the surgery by using ophthalmic ointment (Artificial Tears, Akorn). Alternating Betadine and 70% ethanol swabs were used to sterilize the scalp. Skin was removed from the dorsal surface of the skull. The periosteum was gently removed with a cotton swab and the skull was scrapped with a scalpel. Vetbond (3M) was applied to the wounds at the edge of the skull. A well was made out of dental cement (Metabond, Parkell) around the perimeter with powdered carbon to make the appearance darker. The head post was lowered onto the well and attached with dental cement. Once the dental cement dried, cyanoacrylate glue was applied to the top of the clean and dry skull surface in 4-5 thin layers. Each layer was allowed to dry ∼10 minutes before applying the subsequent layer. Animals were injected subcutaneously with a mixture of Meloxicam (15mg/kg) and enrofloxin (5mg/kg) dissolved in sterile saline (0.9% NaCl, Vetivex). The animals recovered in a cage placed over a heating pad before returning to their home cage. Mice were monitored for at least two days following surgery and injected with Meloxicam (15mg/kg) in sterile saline once per day to manage pain.

### Neuron Reconstruction

#### AAV systemic Injections

Dendrite reconstructions were performed in mice that were injected with a rAAV9-packaged Cre-expressing virus via the superficial temporal vein (STV) of PND1 mouse pups as described previously [50]. Briefly, pups were sedated by covering them with ice for 3 min. STV is visualized using a handheld transilluminator (WeeSight; Respironics, Murrysville, PA, USA), and a pair of standard reading glasses. Virus solution is prepared by 1:50 dilution of stock solution (final ∼ 1 9 1012 GC/mL) in Dulbecco’s phosphate-buffered saline (PBS), also supplemented with 0.001% pluronic-F68. Virus solution (50 nL) was injected using a 100 nL Nanofil syringe attached with a 34 gauge Nanofil beveled needle (World Precision Instruments, Sarasota, FL, USA). A correct injection was verified by noting blanching of the vein. After the injection, pups were returned to the incubator until active and then returned to their dam. Brain slices were cut 13 days after infusions.

#### Histology: ScaleA2 Clearing Solution

For dendritic tracing, P14 animals were deeply anesthetized with pentobarbital (Nembutal) and transcardially perfused with 4% PFA/PBS (wt/vol). Extracted brains were postfixed in 4% PFA/PBS at 4°C for 10 h and cryoprotected in 20% sucrose/PBS (wt/vol) at 4°C for 24 h. Brains were cut on a vibratome (500 μm thickness) collecting both prefrontal and somatosensory cortices. Slices were immediately submerged in Sca*l*eA2 solution in order to clear the tissue. As previously described [51], this urea-based reagent has been reported to be able to render organic tissue transparent after 2 weeks of incubation, without affecting fluorescent signal. Thus, Sca*l*eA2 was implemented in our histological procedure, in order to maximize optical resolution during confocal imaging sessions. After slicing, brain samples (500 µm slice) were placed in 6-well plate and submerged in Sca*l*eA2 solution, at 4_°_C, and gentle shaking. For imaging purpose, tissue was mounted as reported on the published protocol: https://www.labome.com/method/Optical-Clearing-of-Biological-Tissue-with-ScaleA2.html.

Because Sca*l*eA2 clearing-based procedure affects tissue size, we estimate the extent of brain expansion in young (P14, *WT* = 9, *Syngap1^+/-^*= 11) and adult (P60, *WT* = 11, *Syngap1^+/-^* = 9) accounting for *Genotype* and *Age* as main factors. Brain mass weight was measured at different time points: pre-Sca*l*eA2 (T0) and post-Sca*l*eA2 (T24h-T5weeks), and the linear expansion (*E*) was calculated by comparing each post-Sca*l*eA2 measures with the pre-Sca*l*eA2 baseline. The cube root of pre/post Sca*l*eA2 comparisons result in estimating of time-folds linear expansion. Brain mass weight measured in P14 mice revealed that *Syngap1* mutants’ brain was smaller than their control littermates. As a consequence, for comparing morphological parameters of both genotypes, the estimation of the extent of linear expansion promoted by Sca*l*eA2 solution was necessary. Appling the coefficient of expansion (*CoE* = 1/*E*), the measures addressing the total length of dendritic branches were re-scaled accordingly. Because all the confocal sessions were conducted within the weeks post/Sca*l*eA2 third and the fifth, re-scaling *CoE* applied was calculated by averaging each *CoE* from the third to the fifth week during Sca*l*eA2 treatment. The formula [1 – (1/*E*)] was used to determine the percentage of linear expansion in all groups.

#### Imaging and Analysis

When the tissue was transparent (after two weeks), brain slices were mounted in Petri dishes, covered in agarose, and imaged using standard confocal microscopy. Three-dimensional image stacks were collected (x: 2048, y: 2048 pxl; step size 1.00 μm) using confocal microscopy equipped with water immersion objective lens (ULTRA 25x, NA 1.05, Olympus). A computer-based tracing system (Neurolucida360; MicroBrightField) was used to generate three-dimensional neuron tracings that were subsequently visualized and analyzed with NeuroExplorer (MicroBrightField). In order to select a neuron, the following criteria were strictly followed: 1) neuron was selected starting toward the middle of the stack (∼150 μm ± 30 μm) to ensure the accurate reconstruction of an entire dendritic arbor; 2) neuron was distinct from other neurons to allow for identification of branches; 3) neuron was not truncated in some obvious way [27]. Neurons were traced by an experimenter blind to genotype.

#### Spine Density Analysis

Because the clearing tissue method enhanced the quality of our images, spine density was determined using the same set of images previously acquired for the tracing experiment. As previously described [27], ten to fifteen dendritic segments of the SSC-bf L2/3 (20–90 μm in length) at P14 mice were collected and considered for analysis. All measurements were performed by an experimenter blind to the experimental conditions. Pictures were visualized and elaborated with Neurolucida 360 software (MicroBrightField).

### Electron Microscopy Analysis

Young (P14) *Syngap1*+/- (n = 2 mice) and Wildtype (n = 2 mice) were perfused at room temperature with 4% paraformaldehyde + 2% glutaraldehyde in 0.1 M phosphate buffer. Extracted brains were postfixed at 4°C for 2 hours to overnight, then washed in phosphate buffer, and sectioned at 350-400 um. Sections were then washed in 0.1 M cacodylate buffer, then fixed in 2% glutaraldehyde in cacodylate buffer for 15 minutes, washed and then fixed in 1% osmium tetroxide in cacodylate buffer for 30 minutes, washed in buffer and then dehydrated in an alcohol series (with 1% uranyl acetate in the 50% alcohol). Sections were then placed in propylene oxide and finally embedded in epon and polymerized at 64°C. Thin sections were stained with lead citrate and photographed using a JEOL 1010 transmission electron microscope. Layer 1 of SSC-bf was imaged and analyzed.

### Electrophysiology in acute brain slices

Acute coronal brain slices were prepared as previously described from *Zs*green crossed to *Syngap1^+/fl^*mice or conventional *Syngap1* heterozygous mice [30]. *Zs*green mice were injected in the STV of P1 pups as described above, but using 50 µl - virus diluted in PBS to 2.5e+11 vp; (25 ul per side) of pENN.AAV.hSyn.Cre (addgene:#105553). Briefly, P14-P21 mice were sacrificed via cervical dislocation, and their brain was rapidly dissected. 300-350 µm thick slices were cut using a vibratome (7000Smz-2, Campden Instruments), in ice-cold cutting solution containing (in mM): 75 Sucrose, 85 NaCl, 25 D-Glucose, 25 NaHCO_3_, 2.5 KCl, 1.25 NaH_2_PO_4,_ 0.5 CaCl_2_, 4 MgSO_4_ and 0.5 Ascorbic acid (pH7.4, ∼300 mOsm, bubbled with 95% O_2_ and 5% CO_2_). The slices were transferred and warmed to 32°C for 30 minutes in standard artificial cerebrospinal fluid (aCSF), composed of (in mM): 119 NaCl, 2.5 KCl, 26 NaHCO_3_, 2.5 CaCl_2_, 1 NaH_2_PO_4,_ 1.3 MgSO_4_, and 11 D-Glucose (pH7.4, ∼300 mOsm, bubbled with 95% O_2_ and 5% CO_2_). Following this, the slices were maintained in bubbled aCSF at room temperature until transferred to a submerged-type recording chamber for electrophysiological experiments. All experiments were performed at 30-32°C (2-3 mL/mins flow rate), with the experimenter blinded to genotype at the time of recording and throughout the analysis.

Whole-cell patch clamp experiments were conducted from visually identified L2/3 neurons in SSC, M1/M2, PrL/IL and mFC using infrared DIC optics and epifluorescence for *Zs*green (Cre^+^) neurons and regular spiking was confirmed in current clamp mode. Care was taken to alternate between patching Cre^+^ and Cre^-^ neurons within a given brain region to ensure balanced data acquisition for each animal. Recordings were made using borosilicate glass pipettes (3-6 MΩ; 0.6 mm inner diameter; 1.2 mm outer diameter; Harvard Apparatus). All signals were amplified using Multiclamp 700B (Molecular Devices), filtered at 2.4 KHz, digitized (10 KHz), and stored on a personal computer for off-line analysis. Analog to digital conversion was performed using the Digidata 1440A system (Molecular Devices). Data acquisitions and analyses were performed using pClamp 11.4 software packages (Clampex and Clampfit programs; Molecular Devices).

For current and voltage clamp recordings, internal pipette solution contained (in mM): 130 Potassium Gluconate, 5 KCl, 10 HEPES, 0.25 EGTA, 10 phosphocreatine disodium, 0.5 Na-GTP, 4 Mg-ATP and 0.2% Neurobiotin (pH 7.4, ∼290 mOsm). Cells with access resistance >25 MΩ, were unstable (>20 % change in access resistance) or whose resting membrane potential were > −60 mV were discarded from further analysis. Intrinsic properties were recorded at the resting membrane potential for each cell with the following protocols. For voltage clamp protocols, cell capacitance, access and membrane resistance was determined using the membrane test tool in Clampex, while input resistance was measured from small, hyperpolarizing voltage steps (5-10 mV) and was calculated using Ohm’s law. For current clamp experiments, resting membrane potential was calculated from the mean of 1 min recording in gap-free mode. Rheobase was measured from a series short, 2 ms duration, current steps in 50 pA increments. Input/output curves were generated from a series of 800 ms long current steps, from −100-900 pA in 20 or 50 pA increments.

### Organotypic slice culture preparation and electrophysiology

Organotypic slice cultures (OTC) were prepared using the interface method described by [52] with modifications [53]. Briefly, PND 5 (mFC) or PND 10 (SSC) conventional *Syngap1* mice were decapitated and brains submerged in ice-cold slicing solution composed of Hank’s Balanced Salt Solution (+ Calcium Chloride, + Magnesium Chloride) (Gibco), supplemented with Kynurenic acid (3mM) and D-glucose (0.6 % final). mFC or SSC coronal sections (350 μm) were obtained using a vibratome and trimmed to size in ice-cold, bubbled (95 % O_2_, 5% CO_2_) slice solution. A dissecting microscope was used to ensure only brain regions of interest would be cultured. Sections were allowed to rest at 4 °C for 30 minutes and then transferred to culture media for a gentle wash. Culture media was composed of sterile-filtered 25% heat-inactivated horse serum, 50% MEM, 25% HBSS, supplemented with 1 mM Glutamax, 1% D-Glucose, 0.5 mM L-ascorbic acid and 25 U/ml Penicillin/ streptomycin (pH 7.2, 320 mOsm). Sections were mounted onto semiporous membrane inserts (Millicell Cell culture inserts, 0.4 μm pore size, 12 mm diameter) and placed into 24-well plates with 250 μl of culture media, ensuring the section did not make contact with the side of the insert. Slices were moved to a CO_2_-incubator (37 °C with 5% CO2). The day after culturing, slices were treated with vehicle (DMSO, .05%) or drug (PD98059, 10 μm final), with media/drug changes occurring every two days until cultures were used for electrophysiology experiments.

For both SSC and mFC, electrophysiology was started on equivalent PND 17-21; however, due to cultures being prepared at different time points, PND5 for mFC and PND 10 for SSC, they were maintained *in vitro* for 12-16 and 7-11 days, respectively. On the day of electrophysiology experiments, sections were excised from the inserts using a biopsy punch (8 mm diameter) and transferred to warmed (37 °C) aCSF composed of (in mM): 124.06 NaCl, 3 KCl, 25.9 NaHCO_3_, 2.5 CaCl_2_, 1.4 NaH_2_PO_4_, 1 MgSO_4_, and 10 D-Glucose, and bubbled with 95 % O_2_ and 5 % CO_2_ (pH 7.3, ∼320 mOsm). Slices were placed into a submerged-type recording chamber maintained at 32 °C (2-3 mL/min flow rate). All electrophysiology recordings were obtained in a similar manner to that of acute slice recordings except the internal solution osmolarity was ∼310 mOsm.

### Habituation to Head Fixation

Prior to sensory stimulation experiments animals were handled and habituated to head fixation and the insertion of the C2 whisker into a pipette tip attached to a piezo stimulator over the course of four days. On the morning of the first day, animals were handled in their home cage and held by the experimenter for 5 minutes. On the afternoon of the first day, animals were held by the experimenter and allowed to explore the head fixation platform while freely moving. On the second day, animals were briefly handled and then head fixed to the experimental platform. Next the transparent window was cleaned, and the animal was positioned under the macroscope for 10 minutes with imaging and behavior recordings 3-6 minutes during the session. The third day of habituation was similar to the second except the head fixation time was increased to 30 minutes and two recordings were made at 3-6 and 15-18 minutes during the session. Following this habituation to head fixation, animals were briefly anesthetized and all but the C2 whiskers were trimmed. On the final day of habituation, animals were again head fixed and imaged. During this session the head fixation time was increased to 45 minutes, and three recordings were done at 3-6, 15-18 and 42-45 minutes. In addition, prior to imaging the right C2 whisker was inserted into a pipette tip attached to a piezo (as will be done during subsequent whisker stimulation experiments), but no stimulation was given.

### Sensory Stimulations

Each stimulus was repeated pseudorandomly either 25 times (Germline) or 50 times (*Emx1*) during the experimental session such that each stimulus was presented prior to a subsequent repetition. Each stimulus was slightly staggered during the trial by a random delay of 0-250ms. Trials consisted of a baseline 3s, random delay, stimulus presentation, post stimulus time of 6s and an unrecorded 10s intertrial interval.

#### Visual

Visual stimuli were presented on a Rasberry Pi 7” capacitive touch screen LED display (resolution 1024×600, brightness 350cd/m2) placed 13.5cm in front of each mouse triggered by a custom python script. Stimuli were moving black and white gratings angled at 45⁰ or flickering black and white circles generated using PsychoPy (Open Science Tools Ltd.) The gratings and flickering circles were 3.4cm (∼14⁰) wide displayed at full display brightness. The gratings were presented at 100%, 30% or 10% contrast and the flickering circles were alternating between black and white at 3Hz, 10Hz or 30Hz. Stimuli on the screen were centered ∼30⁰ to the right of the nose direction. Gratings were presented for 500ms while flickering stimuli were presented for 1s. Only the grating stimuli were used for *Emx1* experiments.

#### Whisker

The day prior to whisker stimulation, all but the right and left C2 whiskers were trimmed under isoflurane anesthesia (5%, 500mL/min flow for induction, 1.5% 150 mL/min flow for maintenance). If doublet C2 whiskers were present, both were trimmed down to the same length. During the experiment, the right C2 whisker was inserted into a pipette tip attached to a piezoelectric bending actuator controlled by a linear voltage amplifier (Piezo Systems Inc., Woburn, MA) ∼2mm away from the mouse’s whisker pad. The insertion was done using forceps and a micromanipulator. Piezo stimulations were controlled by (BK Precision 4052) waveform generator and triggered by a custom python script. Deflections were either ∼200µm or ∼52µm and calibrated using a laser-based displacement device (LD1610-0.5 Micro-Epsilon, Raleigh, NC). The whisker stimulations were as follows: 5 pulses at 1Hz, 5 pulses at 5Hz, 5 pulses at 10Hz, 1 pulse at 10Hz, 10 pulses at 10Hz, 20 pulses at 10Hz, 20 pulses at 40Hz all with a 200µm deflection and 20 pulses at 40Hz with a 52µm deflection. For *Emx1* experiments, only the 5 pulses at 5Hz and 20 pulses at 40Hz with a 200 µm deflection and the 20 pulses at 40Hz with a 52µm deflection were used

#### Multisensory

All multisensory stimuli were presented the same way as the uni-sensory stimuli except the duration of both stimuli were set to the length of the longest individual one. Stimuli were triggered simultaneously. All multisensory stimuli were presented on a separate experimental day except for the visual-whisker stimuli for the EMX1 animals which were presented along with the whisker uni-sensory stimuli. The stimulus pairs were as follows: *visual-whisker (germline)* – 10% contrast grating/5 pulse at 5Hz 200 µm deflection, 30% contrast grating/5 pulse at 5Hz 200 µm deflection, 10% contrast grating/5 pulse at 10Hz 200 µm deflection and 30% contrast grating/5 pulse at 10Hz 200 µm deflection. *visual-whisker (Emx1)* – 30% contrast grating/5 pulse at 5Hz 200 µm deflection and 10% contrast grating/ 20 pulse at 40Hz 52 µm deflection.

**Widefield Imaging**

Imaging was conducted as described in Couto and Musall et al. 2021 with minor modifications. Mice were head-fixed on a removable platform outside of the imaging box using a head bar and secured to a head bar holder with a screw. Animals were positioned in a transparent acrylic half-tube. The bottom beneath the half-tube was a black acrylic floor with a screen protector (Supershieldz). For imaging sessions that required whisker stimulation or during 45-minute habituation sessions, the C2 whisker was inserted into a pipette tip attached to the end of a piezo mounted onto a x-y-z manipulator. The cured cyanoacrylate glue was cleaned with a micro applicator stick (Patterson Dental, # 70826198) dipped in 70% ethanol prior to imaging. Next, the platform was positioned inside an enclosed sound attenuated imaging box and secured with a magnet. Each animal was then positioned under the macroscope and light shield with the lateral blood vessels in focus.

A custom-made tandem lens macroscope (top lens 105mm DC-Nikkor, Nikon; 85mm 85M-S, Rokinon; PCO Edge5.5 scientific CMOS camera was used for all widefield imaging experiments [49, 54]. For all experiments, images were acquired at 60fps with alternating 405nm (M405L3, Thorlabs) and 470nm (M470L3, Thorlabs) collimated illumination coupled using a dichroic mirror (no. 87–063, Edmund optics; effective 30fps per channel). Light was focused onto the transparent skull using a 495nm long-pass dichroic mirror (T495lpxr, Chroma). GCaMP fluorescence was filtered by a 525nm band-pass filter (no. 86–963, Edmund optics) before entering the camera. To explore the Ca^2+^-dependent signal of GCaMP, the violet excitation was scaled and subtracted from the subsequent frame of blue excitation (see hemodynamic correction below). This subtracted signal was used for all analyses. Following 4 x 4 spatial pixel binning, acquired images were a resolution for 640 x 540 pixels with ∼20µm per pixel. During sensory stimulation experiments, images were acquired only during each 9s trial, and no images were captured during the 10s inter-trial-interval. During habituation, images were acquired continuously for 180s for each of up to 3 imaging windows (see habituation to head fixation).

### Widefield Preprocessing

Widefield image preprocessing was based on methods described in Couto and Musall et al. 2021 with some minor modifications.

#### Motion Correction

To correct for motion of the image, the x and y shifts were estimated using a phase correlation from the first 60 frames [49, 55].

#### Baseline Correction

All widefield imaging data is presented as a baseline zscored ΔF/F signal. For sensory stimulation experiments, ΔF/F was calculated by taking the average 640 x 540-pixel frame of a 1s baseline period before stimulus presentation and subtracting this value each other frame within a trial. This ΔF/F signal was then baseline zscored per trial by subtracting the mean frame within the 0.5s window before stimulus presentation and dividing standard deviation of each pixel within the same window. For spontaneous movements during habituation, ΔF/F was calculated by subtracting the average frame from across a full 3-minute video from each frame of the video. The baseline zscoring for transition movements was done by subtracting the mean of the 5 frames proceeding movement onset and then dividing by the standard deviation of those same frames.

#### Singular Value Decomposition

Singular value decomposition was applied to concatenated trial data from single recording sessions to compute the top 200 highest-variance dimensions. This process leads to the creation of spatial components (U), temporal components (V^T^) and singular values (S). The original data structure can be reconstructed using the product of U and SV^T^. When temporal features were analyzed the SV^T^ component was used to facilitate computations.

#### Hemodynamic Correction

First, both the violet and blue excitation channels were high-pass filtered at 0.1Hz using a second order Butterworth filter. Next, the violet channel was low-pass filtered at 14Hz using a second order Butterworth filter. The violet channel is then rescaled to account for differences in acquisition time and amplitude with the blue channel. Finally, these rescaled violet frames are subtracted from the blue frames to correct for hemodynamic signals [4, 49].

#### Image Alignment

Images were rigidly aligned to a project of the dorsal cortex surface of the Allen Brain Common Coordinate Framework [4, 49]. Using four landmarks from the skull: the left, center and right extent where the anterior cortex meets the olfactory bulbs and the medial point at the base of the retrosplenial cortex.

### Region of Interest Analyses

Once regions were determined (see below), the average signal in the region was quantified by reconstructing the ΔF/F signal from the temporal components SV^T^ and the region of the spatial components corresponding to the region U[region] and averaging spatially across every pixel in the region. The resulting trace was a single average value at each imaging frame for each of the regions.

#### Custom Region of Interest Analysis

Custom regions were selected by visually inspecting macroscope GCaMP6s signal images in the frames following sensory stimulation. Hotspot regions that were consistently visible across animal were selected by drawing a rectangular region over the center of the signal in the earliest visible frame after stimulus presentation. The upper left corner coordinate of this rectangle was used to generate a 10×10 pixel square that was used as the custom region of interest. Regions were drawn for individual animals while the experimenter was blinded to the genotype. For the whisker stimuli the regions labeled included barrel cortex, anterior retrosplenial cortex and secondary motor cortex (termed whisker motor cortex). For the visual stimuli regions included, primary visual cortex, posterior retrosplenial cortex and secondary motor cortex (termed visual motor cortex).

#### Allen CCF Region of Interest Analysis

Images were rigidly aligned to the Allen CCF (described in *image alignment* above). The regions used are from the atlas described in Couto and Musall et al. 2021.

#### Activity Mask Region of Interest Analysis

First, a mask of pixels was determined for each experimental session by taking all the pixels of the control group average signal over 40% (whisker) or 25% (visual) of the max signal from 100-1000ms following stimulus onset. This “activity” mask was then intersected with the regions from the Allen CCF regions described above. To avoid regions with a small number of scattered pixels, only regions that with greater than 500 pixels or accounted for more than 10% of the Allen regions area, were used in subsequent activity mask analysis.

#### Seed-based Correlation Maps

Correlation maps for large movements were calculated for each movement event within a 1.5s window following movement onset. First the covariance of the temporal components SV^T^ was calculated. Next, the covariance between the region and each other pixel as well as the standard deviation was computed. Finally, the division of the covariance by the standard deviation gave the final correlation map.

### Body Movement Recordings

Body movements were recorded with a profile camera (Basler ace 2 a2a1920-160umBAS) placed 7.5cm from the left side of the animal using a 6mm fixed focal length lens (Edmund Optics #33-301). Images were acquired at 60fps and 1920 × 1200 resolution.

### Body Movement Detection

The motion energy within the whisker pad RoI was calculated using FaceMap [6, 56]. The raw motion energy traces were filtered using a one-dimensional gaussian filter to reduce artifacts due to video compression. Filtered data were zscored per trial to generate comparable movement activity across RoIs and animals.

#### Spontaneous Movements

spontaneous movements of the animal were identified by using a two-pass method. First, motion energy above an empirically determined threshold of 0.7 *z*-score units were flagged as movements. Second, a rolling threshold was calculated using the sum of the root mean squared error and the rolling mean of a 20-frame window. All the remaining frames with zscored movement above this threshold line were also considered movements.

#### Movement Pre and Post Stimulus

Trials from sensory stimulation experiments were grouped based on whether movement was detected in a 0.25s window before or after stimulus onset. Movements within these windows used the same two-pass method as spontaneous movements but also added an additional two passes for further refinement. The first additional pass was to include trials as movement based on the area under the curve (AUC) of the difference between the zscored motion energy rolling threshold. Through visual inspection, we empirically determined trials with an AUC value above 15 were considered to have movement. Trials which had a great deal of movement led to a relatively low zscore. Therefore, to capture these trials, we added a final pass that compared a zscore calculated from all the motion energy traces stitched together across animals and trials (stitched zscore). We then flagged trials that had a stitched zscore greater than the mean of the trials that had been flagged in the previous three passes.

### Body Movement Quantification and Grouping

#### Breathing Artifacts

only movements above 83ms were included for subsequent grouping and analysis. Visual inspection of the motion energy traces along with paired behavioral videos showed a low amplitude rhythmic signal which aligned with the breathing of the mice. While generally our threshold cutoff removed breathing, on occasion this breathing signal could be miscounted as spontaneous movement for a few frames (<83ms). To clearly separate breathing from spontaneous movement events, we included only movements that lasted for at least 100ms.

#### Movement Events

A movement event was defined by an onset (non-movement frame to movement frame) and offset (movement frame to non-movement frame). Incomplete movements missing either an onset (start before recording) or offset (end after the recording) were discarded from subsequent analyses.

#### Movement Windows

Movements that happened close in time (<0.5s) were clustered together into movement windows. The first movement within a movement window was considered a movement transition (**Fig 3B**) with a subset of these used in subsequent analyses (see movement transition grouping below).

#### Grouping Movement Transitions

Movement transitions were grouped for two reasons. First, to compare relatively similar magnitude movements between genotypes (**Fig. 3D,G,J and Fig. 4A,B,C**) and second to understand how neuronal activity scaled in relation to movement. To group movement transitions into small, mid and large categories we relied on AUC and duration and empirically determined cutoffs by inspecting the movement traces. Small movements were defined by an AUC less than 7 (motion energy units) and a duration less than 150ms. Mid movements were defined with an AUC >= 7 and <= 65 as well as a duration <= 0.5s. Large movements were defined by an AUC > 65 and duration <=1s. These parameters selected movements with clear onset and offset rather than more complex dynamics to which may be due to multipart movements happening in sequence (for example see large movement trace in **Fig 7B**).

#### K-Means Clustering of Movements

All movement events were clustered using k-means clustering. 3 clusters were determined using the elbow method from a plot of inertia by cluster number (**Fig 7A**). K-means clustering was implemented using Python 3.13.5 Scikit-Learn library. To make results reproducible, a deterministic random start seed was selected by running the clustering using 1000 different random state values and selecting the one that produced the lowest inertia. Spontaneous movements from germline *Syngap1* lines and Emx1 *Syngap1* floxed lines were clustered separately, but the clustering led to similar groupings of the movements (**Fig 7A**).

### Statistics

#### Neuronal Tracing and EM

To determine the distribution and the homogeneity of variance all datasets were explored by using Levene’s and Kolmogorov-Smirnov test, respectively. For the neuronal morphology analysis, data are expressed as the mean ± SEM. Analysis of layer 2/3 and 5 neurons were performed by using unpaired t-Test, assessing the contribution of total length, # of nodes and the spine density, in revealing differences between the groups tested in our experiments. Statistical comparisons between regions and genotypes were assessed with two-way ANOVAs and follow up Bonferroni-corrected multiple comparison t-Tests.

#### Macroscope Traces

Data are represented as mean ± SEM with each trial treated as an independent observation. Normality was assessed using D’Agostino and Pearson’s tests. Statistical comparisons are from aligned rank transform nonparametric factorial ANOVA’s with follow up Bonferroni corrected Mann_Whitney U tests for multiple comparisons. Statistical comparisons were conducted using Python statsmodels and pyartool packages.

#### Electrophysiology

All data are represented by mean ± SEM, with neurons treated as independent observations. For statistical comparisons, 2-way ANOVA’s with Tukey’s or Dunnett’s multiple comparisons test were used throughout, except for input/output curves where a mixed-model with repeated measures was used with Tukey’s multiple comparisons test. All statistical comparisons for the electrophysiology data were conducted using Graphpad Prism 10 software.

#### Movement Cumulative Probabilities

Cumulative probability distributions were compared between genotypes for log-transformed peak amplitude or duration across small, mid and large movement clusters. Statistical comparisons were from two sample Kolmogorov-Smirnov tests.

---

**Figure S1 (extended data).**
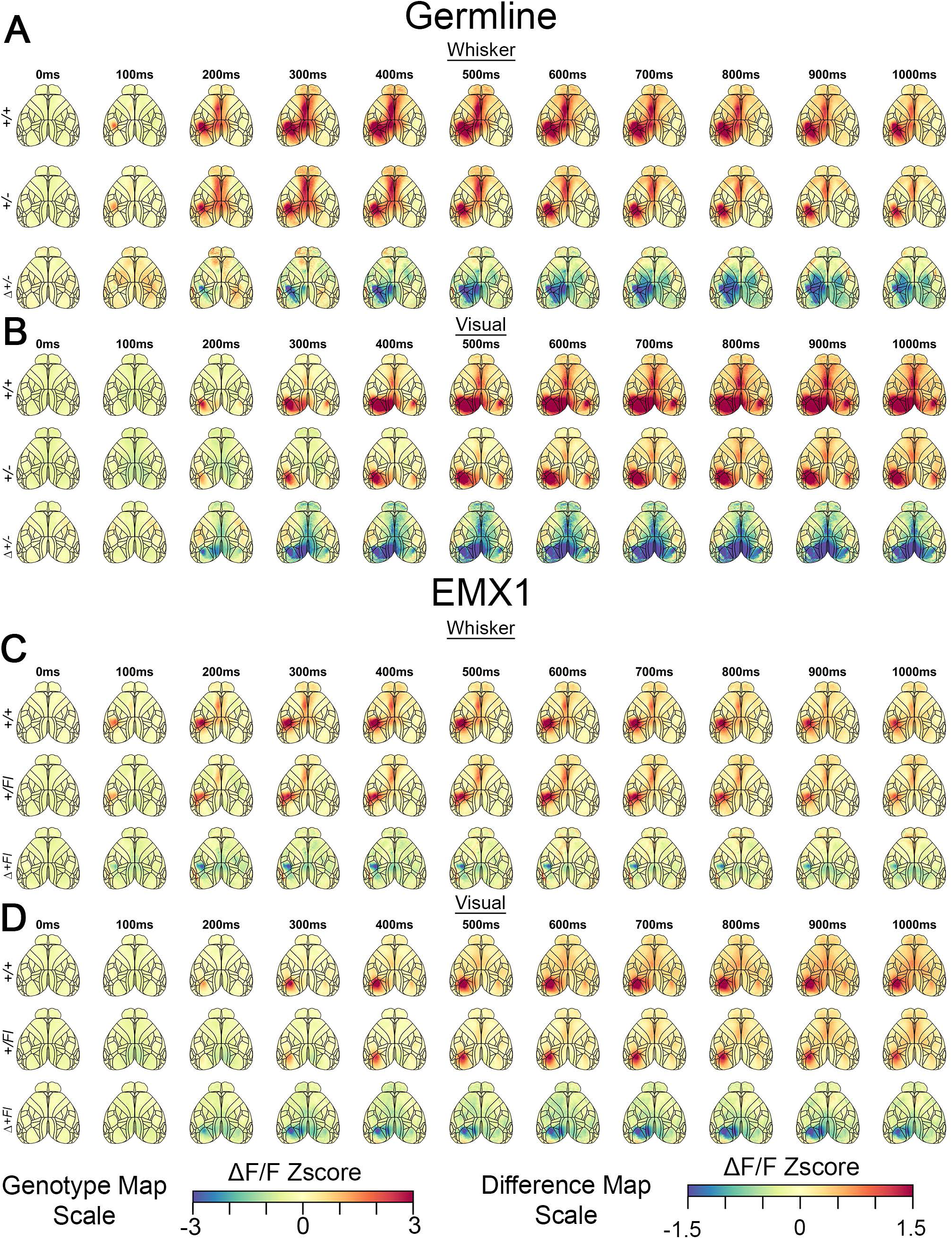
Signal amplitude changes across time in dorsal cortex from different Syngap1 lines in response to either whisker or visual stimuli. **(A-D)** Time series maps for either whisker or visual stimuli in germline and EMX1-Syngap1 strains. Maps were generated at specific time points after stimulus onset for each genotype and a third map was generated reflecting the regional differences by subtracting wildtype activity from mutants.

**Figure S2 (extended data).**
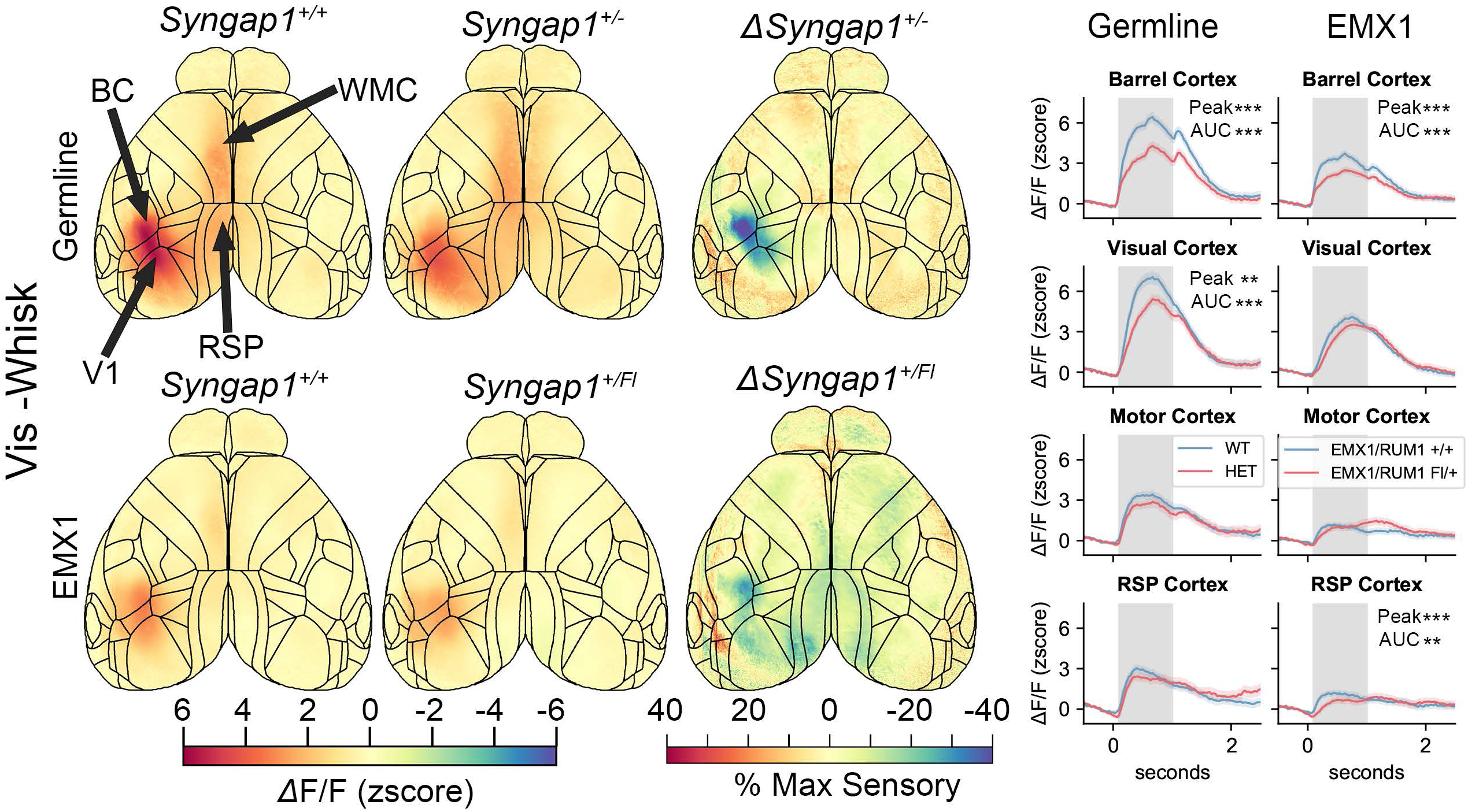
Time-compressed maps of dorsal cortex GCaMP6 signal amplitude in response to multi-modal stimuli in two Syngap1 mutant lines. Maps were generated as shown in main Figure 1 for germline and Emx1-Syngap1 strains but here signals were generated in response to combined visual-whisker stimuli.

**Figure S3 (extended data).**
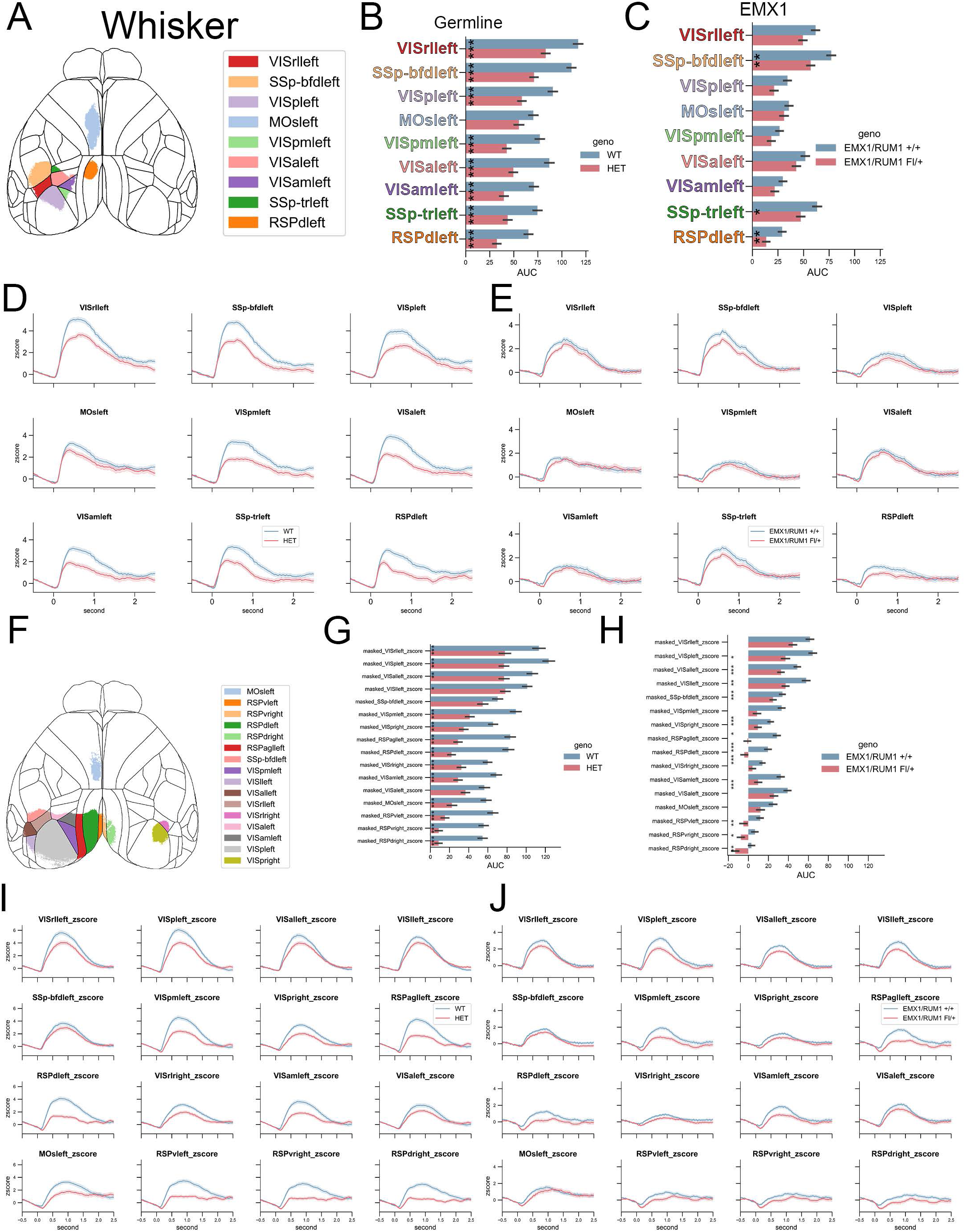
Regional analysis of GCaMP6 signals across dorsal cortex using a masking approach. Masks were created based on WT signals originating from whisker stimuli **(A-E)** (see methods) or visual stimuli **(F-J)** in both Syngap1 mutant strains. Time series for regions within these masked areas are shown.

**Figure S4 (extended data).**
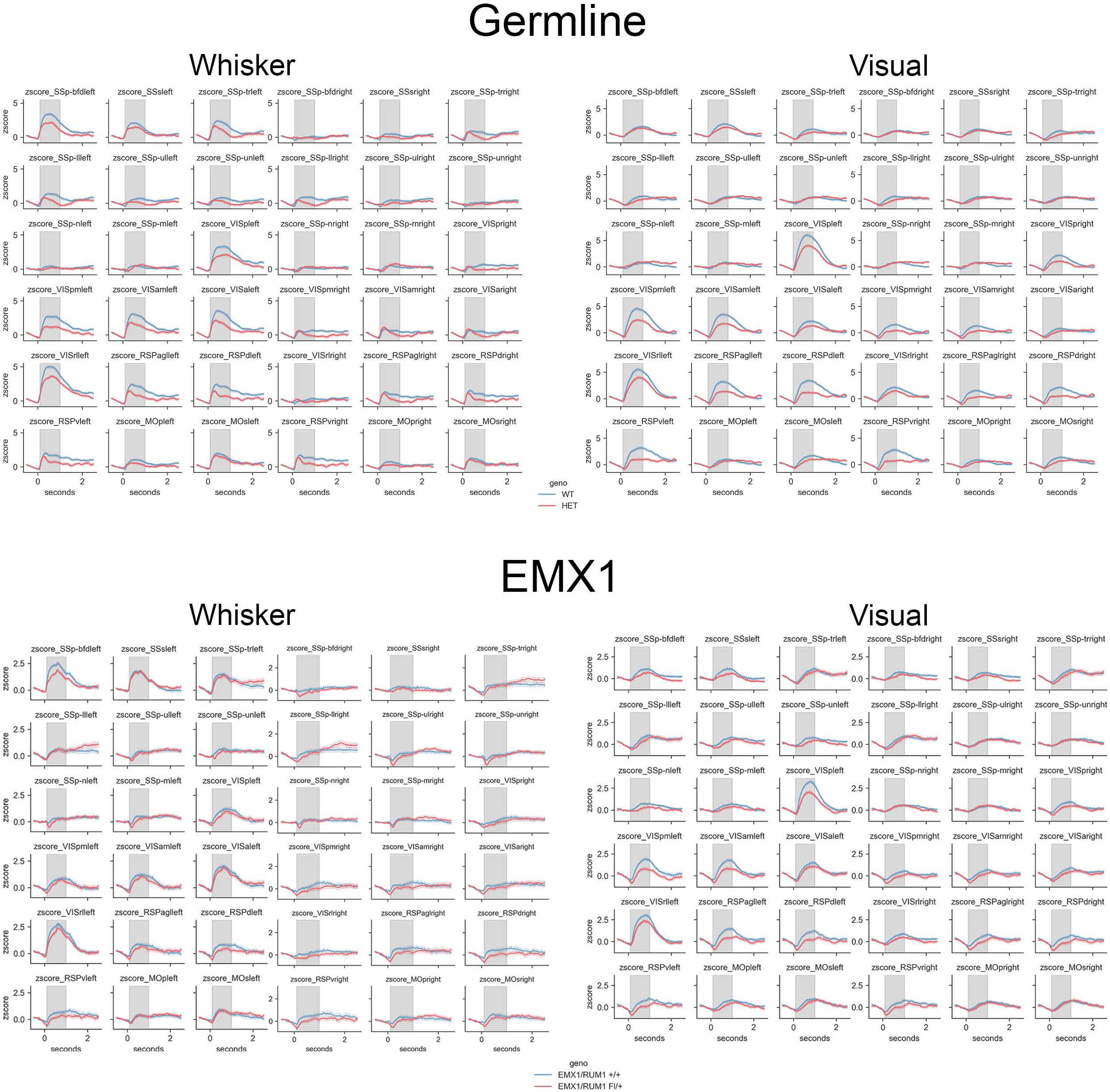
Regional analysis of GCaMp6 signals across dorsal cortex using the Allen Common Coordinate Framework (CCF). Dorsal cortex was parsed into CCF "regions", which were used as ROls to obtain time series for GCaMP6 signals in both Syngap1 strains for either whisker or visual stimuli.

**Figure S5 (extended data).**
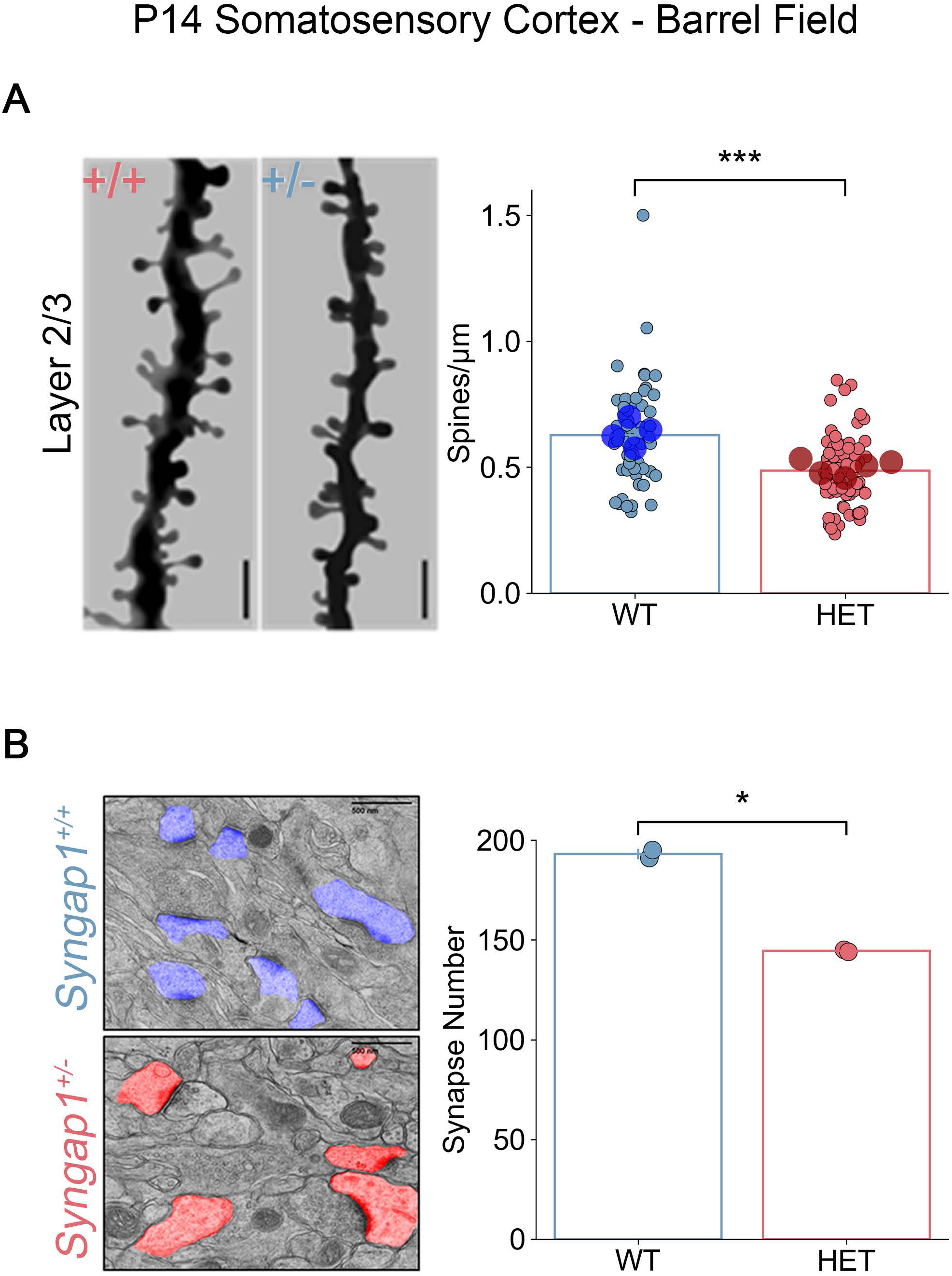
Synapse density measurements in L2/3 of somatosensory cortex from postnatal day 14 germline Syngap1 mice. **(A)** Dendritic spine density of TD-tomato-expressing neurons in **L2/3** of somatosensory cortex. Scale bar represents 5µm **(B)** Dendritic spine synapse in **L2/3** of somatosensory cortex in electron micrographs were identified, labeled, and counted. Scale bar represents 500nm.

**Supplemental Video 1.** (See attached file). Videos examples of small (top), mid (middle) and large (bottom) spontaneous movements from germline control mice with corresponding motion energy traces and widefield calcium activity both plotted on equal scales. Videos have been slowed to 25% playback speed from the original 60 FPS. (left) Video of the mouse during a movement event. The red box over the whisker pad is the region of interest (Roi) for the motion energy calculation from the video. (center) The motion energy trace from the Roi with the vertical black line indicating the onset of the flagged movement event. (right) ΔF/F GCaMP6s signal from widefield calcium imaging with deeper reds indicating more positive regions of activity and deeper blues indicating more negative regions.

**Figure S6 (extended data).**
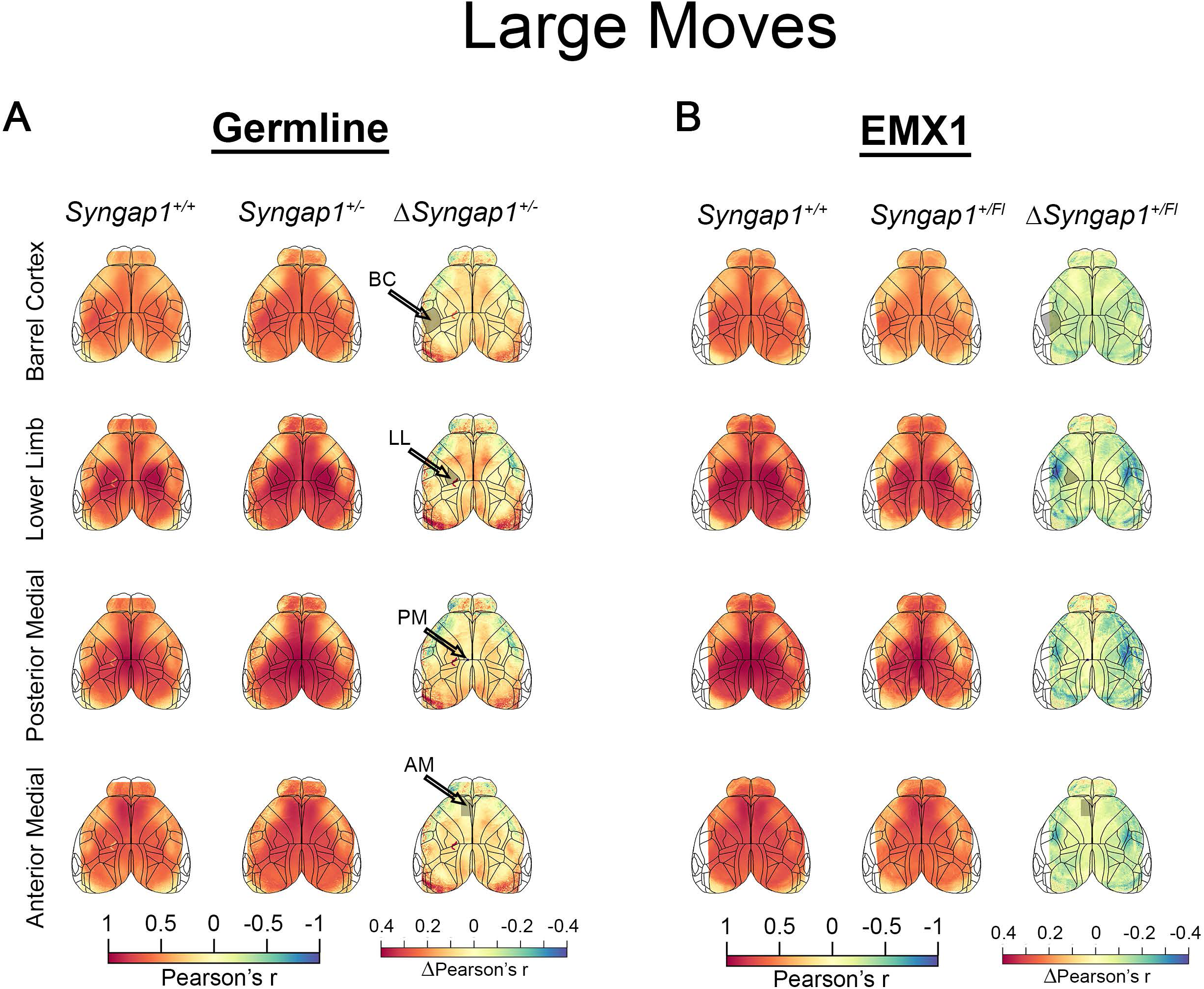
Region-seeded co-activation maps from germline and Emx1-restricted Syngap1 mutants. **(A)** Region-seeded co-activation analyses during large movement-related state transitions in wild-type mice and germline Syngap1 heterozygous mice. Time-compressed co-activation maps are shown for four representative seed locations (barrel cortex, lower-limb somatosensory cortex, posterior medial cortex, and anterior medial cortex). For each seed, difference maps are shown in which co-activation in wild-type mice is subtracted from Syngap1 heterozygous co-activation, revealing a mixture of increased and decreased co-activation across dorsal cortex. (B) Region-seeded co-activation analyses during large movement-related state transitions in Emx1-Cre Syngap1 heterozygous mice, shown as in (A). Across seed locations, Emx1-restricted mutants exhibit predominantly reduced co-activation relative to wild-type controls, indicating a distinct pattern of altered inter-regional co-activation compared to germline mutants.

**Figure S7 (extended data).**
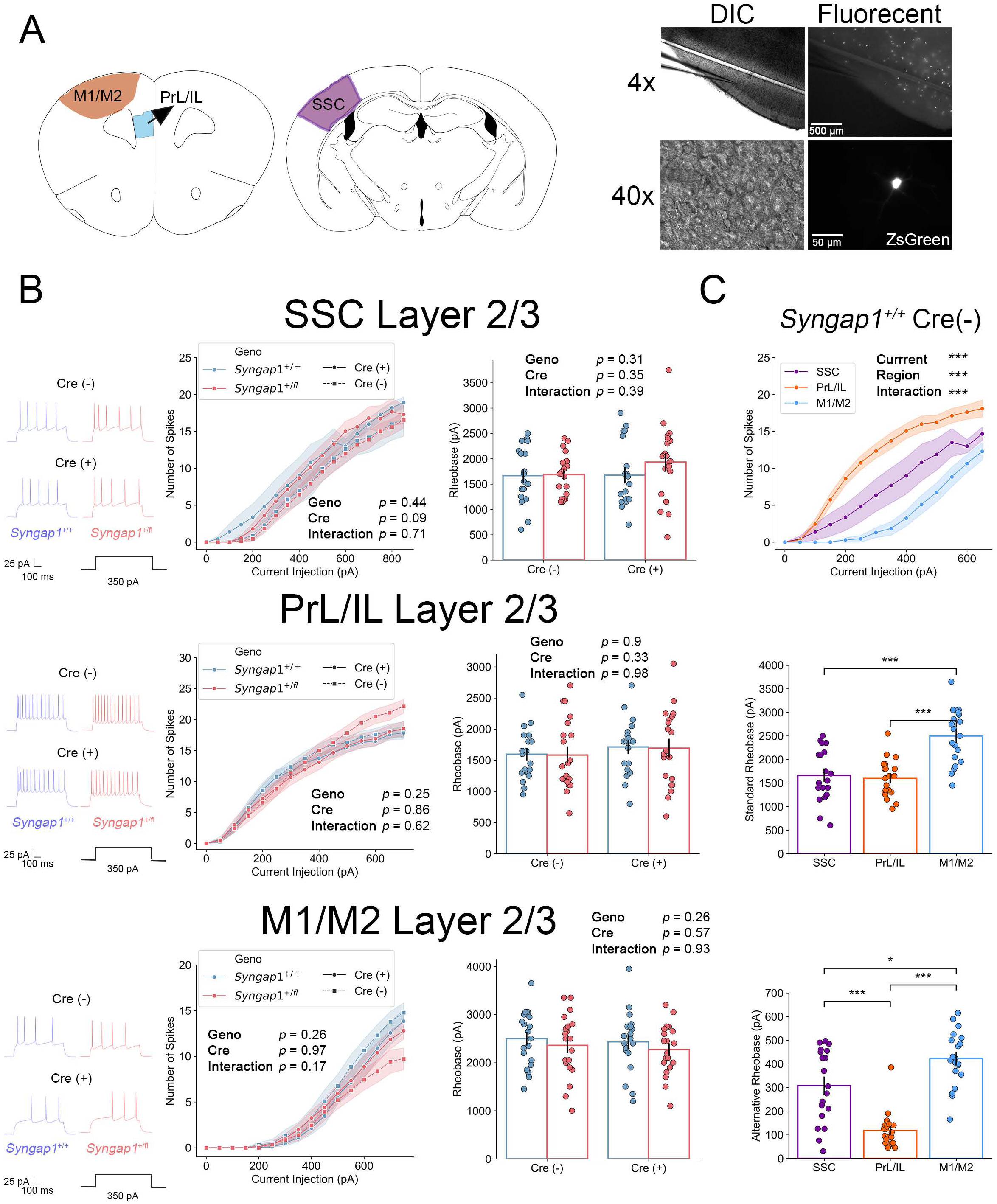
Examining the effect of sparse Cre on intrinsic excitability across cortical L2/3 IT neuron populations in Syngap1 conditional mice. (A) Map noting the L2/3 IT populations examined and the relative number of neurons demonstrating Cre activity in acute slices cut from PND14-21 animals. (B) Measures on intrinsic excitably in each L2/3 IT population noted in (A). For each region examined, both genotype (Syngap1+/+ and Syngap1+/fl) and Cre activity state [(fluorescent, Cre(+)) or (non-fluorescent, Cre(-)] were plotted. Two-factor ANOVA. (C) Syngap1 +/+; Cre(-) populations from each region were plotted to fingerprint the developmental excitability landscape at this developmental time point. RM-ANOVA. All recordings were done by the same blinded investigator using identical experimental approaches.

**Supplemental Table 1 - Comprehensive Statistics Table.** (See attached file). The details about the statistical tests, sample sizes and outputs from all the statistical tests referenced in the manuscript text or figures. Each sheet contains the statistics related to an individual figure or supplemental figure.

